# Specific gut pathobionts escape antibody coating and are enriched during flares in patients with severe Crohn’s disease

**DOI:** 10.1101/2023.06.30.545711

**Authors:** Carsten Eriksen, Niels B. Danneskiold-Samsøe, Janne M. Moll, Pernille N. Myers, Pi W. Bondegaard, Simone Vejrum, Tine B. Hansen, Lisbeth B. Rosholm, Philipp Rausch, Kristine H. Allin, Tine Jess, Karsten Kristiansen, John Penders, Daisy MAE. Jonkers, Susanne Brix

## Abstract

Patients with Crohn’s disease (CD) exhibit great heterogeneity in disease presentation and treatment responses, where distinct gut microbiota-host interplays may play part in the yet unresolved disease etiology. We here characterized absolute and relative single and multi-coating of gut bacteria with immunoglobulin (Ig)A, IgG1, IgG2, IgG3 and IgG4 in CD patients and healthy controls. Patients with severe disease exhibited distinctly higher gut bacterial IgG2-coating. IgG2-coated bacteria included both known pathogenic and non-pathogenic bacteria that co-existed in communities with two non-coated gut pathobionts *Campylobacter* and *Mannheimia*. These latter two exhibited low prevalence, rarely coincided, and were strongly enriched during disease flares in CD patients across independent and geographically distant cohorts. Since antibody-coating of gut pathobionts diminishes epithelial invasion and inflammatory processes, escape from coating by specific gut pathobionts may be a mechanism related to disease flares in the subgroup of CD patients with severe disease.

---

Crohn’s disease (CD) is a chronic and relapsing inflammatory bowel disease (IBD) with great heterogeneity in disease presentation, progression, and treatment responses^1^. There is currently no curative treatment for CD, making monitoring of mucosal inflammation crucial in order to limit disease progression and complications. It is well substantiated that gut bacteria may induce immune activation during the course of CD as demonstrated by the consistent associations between certain gut bacteria and CD disease etiology^2^. Still, we have limited evidence that specific bacteria are consistent disease drivers among patients. Classically, the immunopathogenesis of CD has been described to be driven by uncontrolled Type 1 and Type 17/3 immune reactions with IL-12p70 and IFN-ɣ^3^, and IL-17A, IL-21, IL-22 and IL-23^4^ as the major disease-associated cytokines. These immune reactions may result in distinct effector responses in the form of distinct immunoglobulin (Ig) production, but still the involvement of different IgG isotypes in disease protection or exacerbation in CD remains unresolved. Cytokines associated with Type 1-immunity have previously been described to prime the production of the IgG2 subtype^5, 6^, while the influence of Type 17/3-immune related cytokines on class-switching is less well documented. There are presently few reports on a role for IgG in the anti-microbial barrier defense in the intestine, although IgG is recognized as the major antibody class of circulating blood in bacterial infections^7^. A recent study identified very limited fractions (0.16%) of gut bacteria to be coated with IgG^8^, despite serum-derived IgG has been shown to hold the capacity to bind various gut bacteria^8^. The latter emphasizes that IgG indeed may be induced in the gut mucosa, from where they may be transported to the gut lumen via binding to the neonatal Fc receptor^9^. IgG responses are less promiscuous than those associated with IgA^10^ due to their requirement for T-cell help in isotype switching to IgG, resulting in increased antibody affinity towards targeted antigens^11^.

We here hypothesize that gut bacterial Ig-coating patterns, including the IgG isotypes IgG1, IgG2, IgG3 and IgG4 as well as the notoriously present mucosal IgA, may be used to define underlying immune-mediated processes in CD, thereby helping in differentiating disease endophenotypes. We first characterized single- and multi-coating of gut bacteria with IgA and the four IgG isotypes (in quantitative and relative numbers) in 20 healthy controls and 60 CD patients from the IBD South Limburg (IBDSL) cohort, a population-based inception cohort from the South Limburg area of the Netherlands^12^. The CD patients were all sampled during remission and active disease stages. We identified IgG2-coating in patients with severe CD and increased gut bacterial IgG2-coating during active disease. Taxa comparisons of sorted and sequenced IgG2-coated bacteria versus bulk stool sequenced bacteria led to identification of IgG2-coated bacteria that co-occur with two non-coated gut pathobionts in patients with severe CD during disease flares. The two non-coated gut pathobionts were also enriched during active disease flares in a non-related American cohort of 297 CD patients. Thus, IgG2-coated gut bacteria were identified in patients with severe CD, where they co-occur with distinct non-coated gut pathobionts that appeared during flares, hence pointing to an immunologically uncontrolled presence of certain gut pathobionts during CD flares.

## Results

### IgG2 gut bacterial coating associates with CD severity

Previous studies have reported on the binding of IgA^13, 14^ and total IgG^15^ to gut bacteria in humans. However, bacterial coating with the four IgG isotypes has not been thoroughly studied, although they may be involved in barrier protection in the intestine. To examine the dynamics of IgA and IgG1-4 bacterial coating in healthy individuals and patients with varying severity of CD, we determined the levels of coated gut bacteria in 60 CD patients and 20 healthy individuals (**Supplementary Table 1** for cohort statistics), as relative and actual numbers of coated bacteria per gram of stool, using multi-parametric flow cytometry (**Figure 1a, Supplementary Figure 1**). The median relative abundance of 3.75% IgA-coated gut bacteria in healthy individuals (**Figure 1b**, **Table 1**) was consistent with the average relative abundance reported by Fadlallah *et al.* (2018)^8^ in healthy Europeans. An average of 1.7-fold more IgA-coated bacteria were identified in CD patients compared to healthy individuals (**Figure 1b**, **Table 1**, *P* = 0.024), which is in line with findings in Palm et al.^14^.

**Figure 1.**
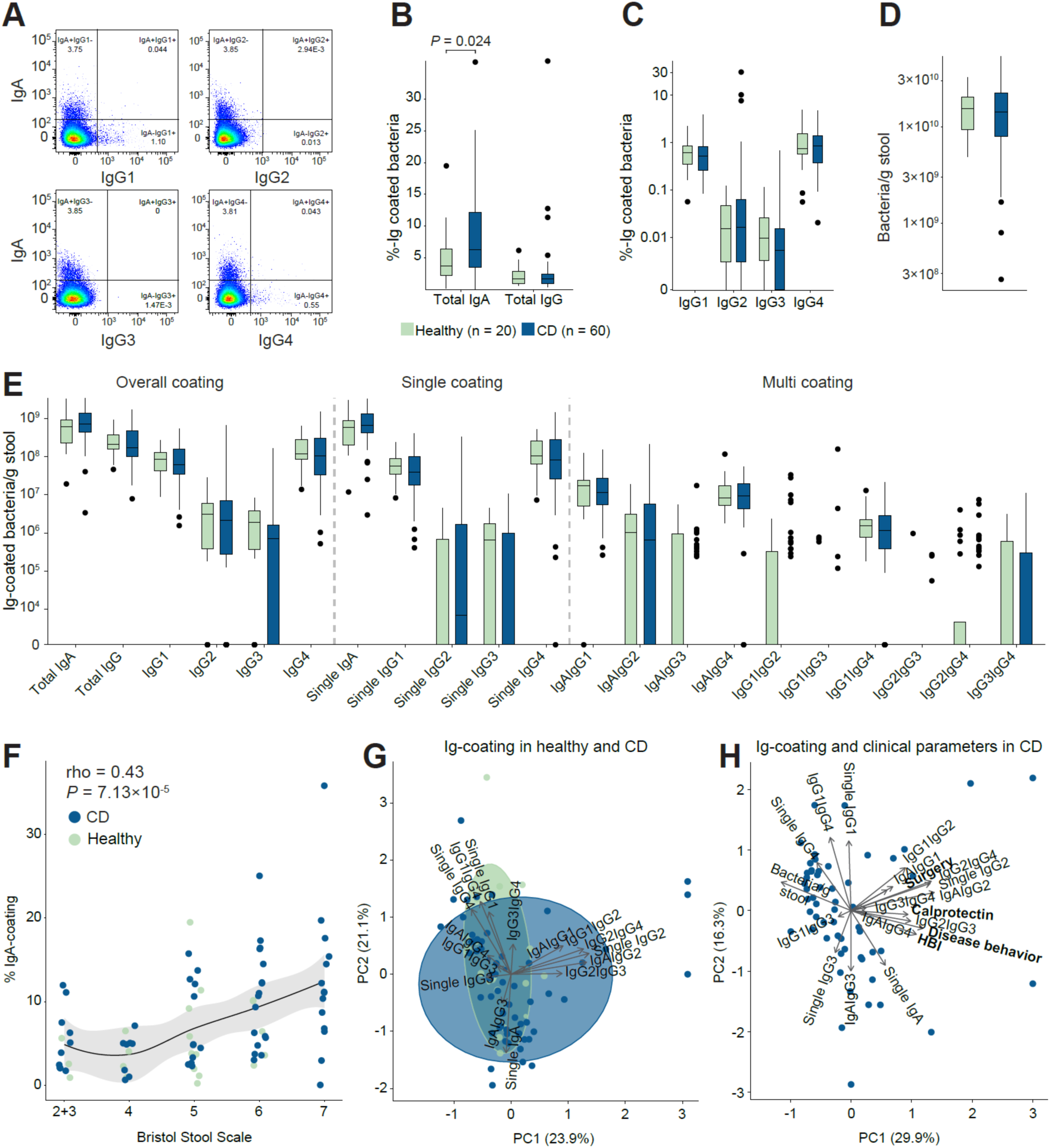
Immunoglobulin-coated bacteria in CD patients and healthy individuals. a) Representative plots of the multi-parametric flow cytometry-based analysis of IgA-, IgG1-, IgG2-, IgG3-, and IgG4 coating of gut bacteria in healthy individuals and CD patients to determine (b) the relative Ig-coating of total IgA and total IgG (sum of IgG1-4), and (c) individual IgG’s. d) Total number of bacteria/g stool based on flow cytometry analysis. e) Quantity of gut bacteria/g stool coated with the different antibodies alone (single) or in combination (multi). The overall coating is the sum of single and multi-coating for the respective antibodies. f) Bristol stool scale vs. % IgA coating of gut bacteria. Spearman’s rho statistics was used for correlations. The line depicts a local polynomial regression fit, and the shaded area is the 95% confidence interval. g) Principal component analysis of Ig-coating frequencies in healthy individuals and CD patients. Dots and the confidence ellipses of the variance within each group are represented by the green color for healthy individuals and blue for CD patients. h) Gut bacterial Ig-coating vs. clinical parameters (calprotectin, HBI, disease behavior, and GI surgery) in CD patients. In all plots: healthy individuals, n=20; CD patients, n=60. Statistical analyses were based on Wilcoxon rank-sum test (b-e) for group comparisons shown as boxplots where center lines indicate the median and the box limits indicate the quartiles. Whiskers extend to the data points within 1.58x the interquartile range, and outliers are shown as individual dots where center lines indicate the median and the box limits indicate the quartiles.

**Table 1.**
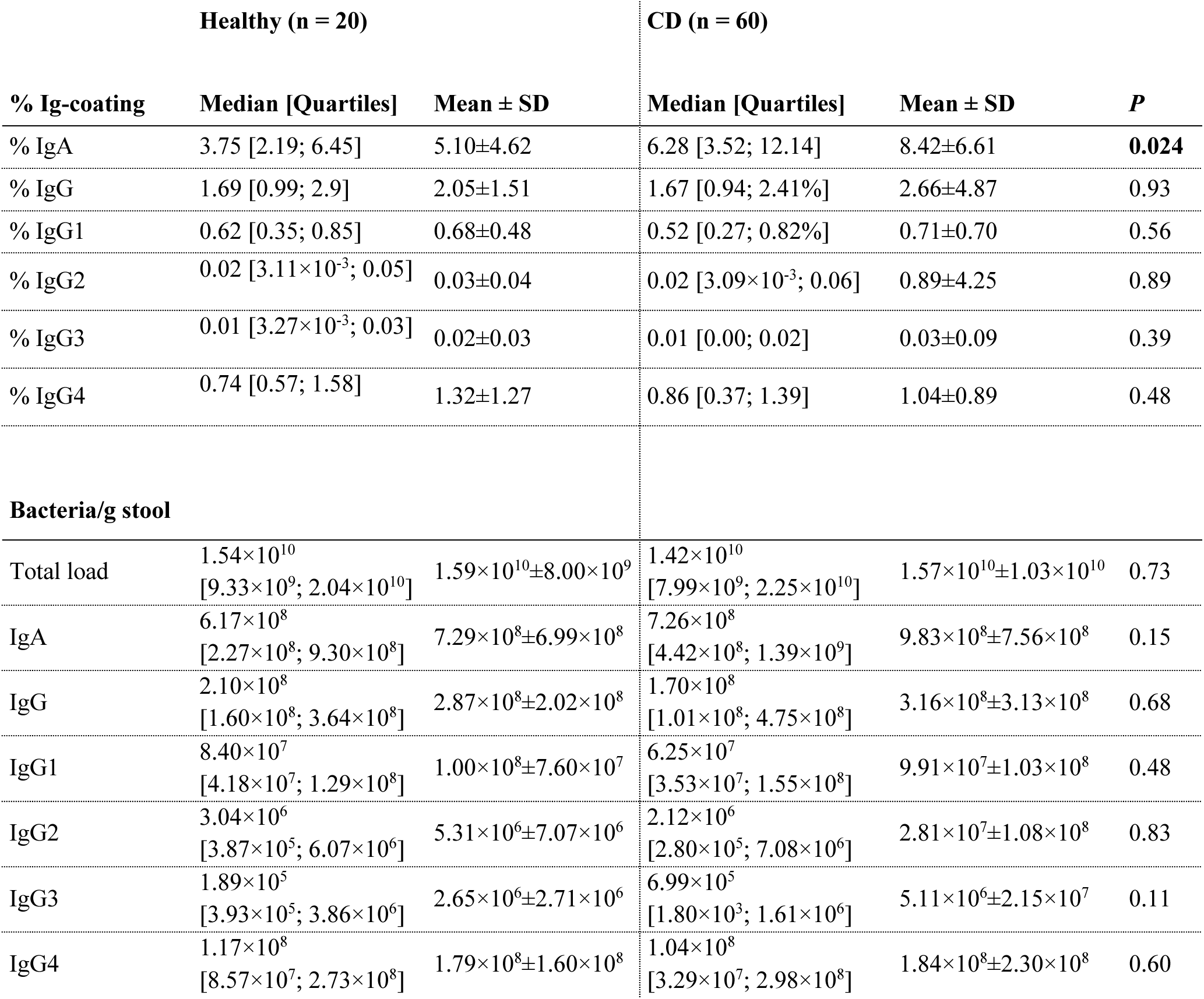
Relative and quantitative Ig-coating of gut bacteria.

The relative bacterial coating with total IgG was only 2.2-fold lower than that with IgA, and did not differ between CD patients and healthy individuals (**Figure 1b**, **Table 1**), thus pointing to substantial gut bacterial IgG-coating independent on disease. The IgG isotypes coated gut bacteria with varying frequency where consistently high coating levels were seen for IgG1 and IgG4, while coating with IgG2 and IgG3 was relatively rare, and no significant differences were identified in the overall IgG-coating between healthy individuals and CD patients (**Figure 1c**, **Table 1**).

Flow cytometry-based counting of bacteria was used to determine the total bacterial load per gram of stool thereby enabling calculation of the actual number of coated bacteria per gram of stool. The bacterial load between healthy individuals and CD patients was not significantly different in this cohort (**Figure 1d**, **Table 1**). Likewise, when examining single and multi-Ig coated gut bacteria frequencies, we found no differences in the actual number of gut bacteria with single or double Ig-coating between healthy individuals and CD patients (**Figure 1e**). More than 96% of the coated gut bacteria were found to be single-coated with IgA and IgG1-IgG4 in both healthy individuals and CD patients (**Supplementary Table 2**). For IgG2, we identified only minute levels of single IgG2-coating of bacteria in the healthy and CD gut; rather, most IgG2 coating co-occurred with IgA (**Figure 1e**).

The % of IgA-coated bacteria was found to correlate positively with an increasing Bristol Stool Scale value (**Figure 1f**, rho = 0.43, *P* = 7.13×10^-5^), a qualitative measure for fewer gut bacteria, reduced stool consistency and faster stool transit time^16^, and also correlated inversely with increasing bacterial load (bacteria/g stool, **Supplementary Figure 2**, rho = -0.46, *P* = 2.48×10^-5^). Hence illustrating that individuals with fewer gut bacteria displayed a higher relative IgA coating of bacteria.

To investigate the dynamics between the different gut bacterial Ig-coating patterns, the single and multi-coating patterns of all individual were visualized in a PCA. Some patients were identified to separate from the rest due to their IgG2 single- and multi-coating patterns (**Figure 1g**; arrows to the right). When we correlated clinical parameters with Ig-coating patterns in CD patients, we found single- and multi-coating with IgG2 to associate with several features related to disease severity (Fecal calprotectin, surgery, Montreal classification of disease behavior (B1-3), and Harvey-Bradshaw Index (HBI)) (**Figure 1h**).

### Gut bacterial IgG2-coating is enhanced in CD patients with active disease and high IgA-coating

We next stratified the cohort based on IgG2-coating tertiles, resulting in three IgG2-coating phenotypes (IgG2-low (IgG2-lo; 0.0% [0.00%; 0.003%] (median [25^th^; 75^th^ Quartile])), IgG2-intermediate (IgG2-int; 0.02% [0.007%; 0.03%]) and IgG2-hi (IgG2-hi; 0.24% [0.09%; 0.58%]) (**Figure 2a**). Healthy individuals were only represented within the IgG2-lo group, while CD patients in IgG2-hi displayed higher numbers of IgG2-coated bacteria/g stool during active disease vs. remission (**Figure 2b**, *P* = 0.028). It is noteworthy that CD patients with active disease can hold any of the three IgG2-coating levels (IgG2-lo, N=8, IgG2-int, N=11, IgG2-hi, N=8), stressing that bacterial IgG2-coating is not a generic marker of active disease, although it links to CD severity.

**Figure 2.**
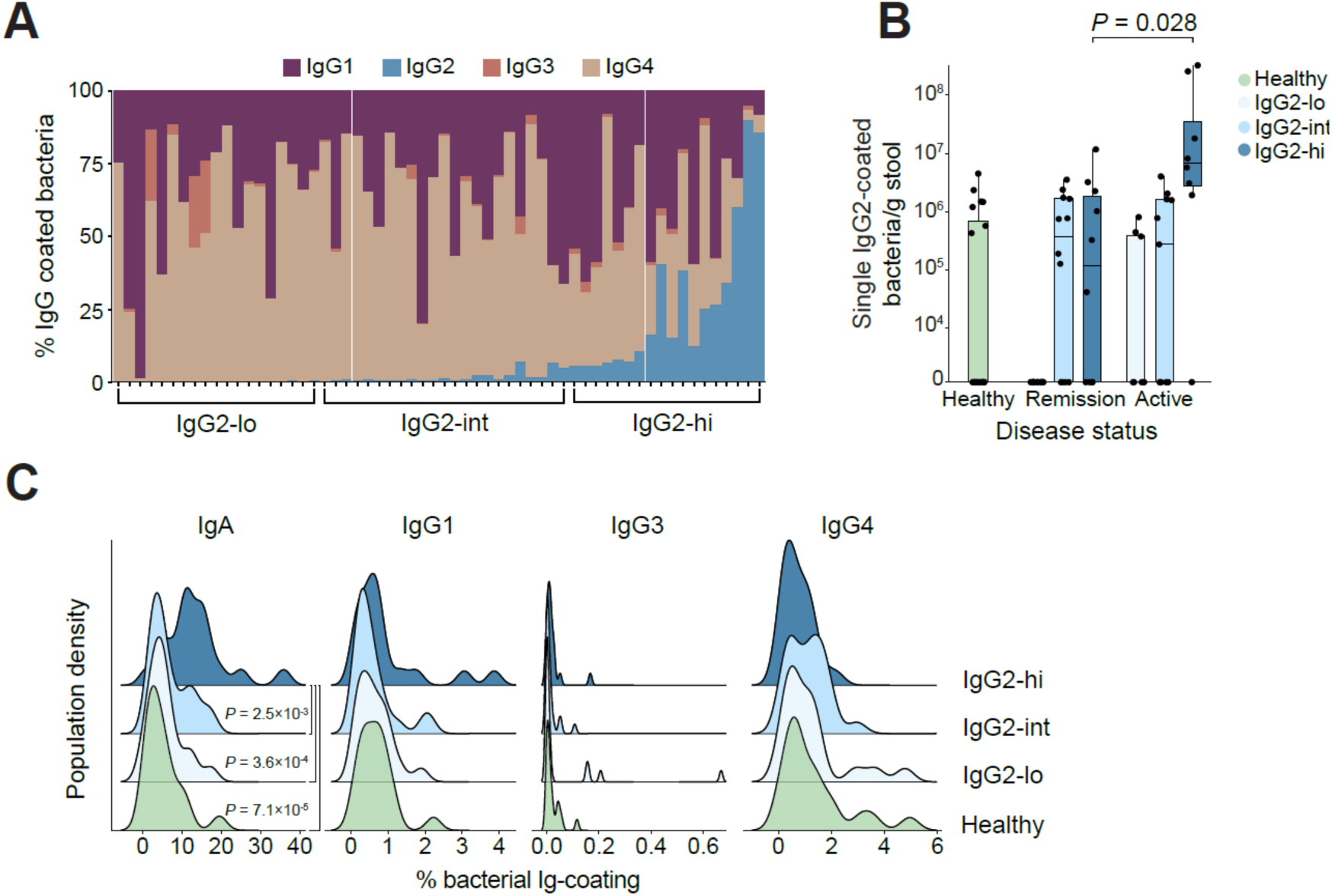
CD patients exhibit differential gut bacterial IgG2-coating during active disease. a) Subgrouping of CD patients based on tertiles of gut bacterial IgG2-coating levels. b) Load of single IgG2-coated bacteria/g stool in healthy controls (n=20) and CD patients in remission (n=33) or with active disease (n=27) for each IgG2-coating subgroup. Center lines of box plots indicate the median and the box limits indicate the quartiles. Whiskers extend to the data points within 1.58x the interquartile range. Dots represent the level within each individual. c) Joyplot illustrating population densities of IgA and IgG1, IgG3 and IgG4 in healthy individuals and CD patients stratified on the IgG2-coating subgroup. Statistical analyses were based on Wilcoxon rank-sum test (b-c) for group comparisons.

IgA-coating in IgG2-hi (11.68% [10.43%; 15.66%]; median [25^th^; 75^th^ Quartile]) was significantly enhanced compared to IgG2-lo (4.89% [2.72%; 6.7%], *P* = 3.6×10^-4^), IgG2-int (4.82% [3.16%; 10.23%], *P* = 2.5×10^-3^) and healthy controls (3.75% [2.19%; 6.45%], *P* = 7.1×10^-5^), while concurrent IgG1-, IgG3- and IgG4-coating did not differ significantly between healthy and IgG2 subgroups (**Figure 2c**).

### Twenty-five indicator genera characterize the gut microbiota in IgG2-hi CD patients

The load of gut bacteria in CD patients, defined as bacteria per gram of stool, was found to strongly associate with the bacterial α-diversity calculated using the Shannon index (**Supplementary Figure 3a**, rho = 0.747, *P* < 2.2×10^-16^), indicating that individuals with a low Shannon index have a lower bacterial load. Patients with IgG2-hi bacterial coating versus IgG2-lo and IgG2-int displayed a significantly lower Shannon index (**Figure 3a**, IgG2-hi vs IgG2-lo, *P* = 0.015 and IgG2-hi vs IgG2-int, *P* = 0.010), as well as reduced bacterial richness (**Figure 3a**, IgG2-hi vs IgG2-lo, *P* = 0.013 and IgG2-hi vs IgG2-int, *P* = 0.013).

**Figure 3.**
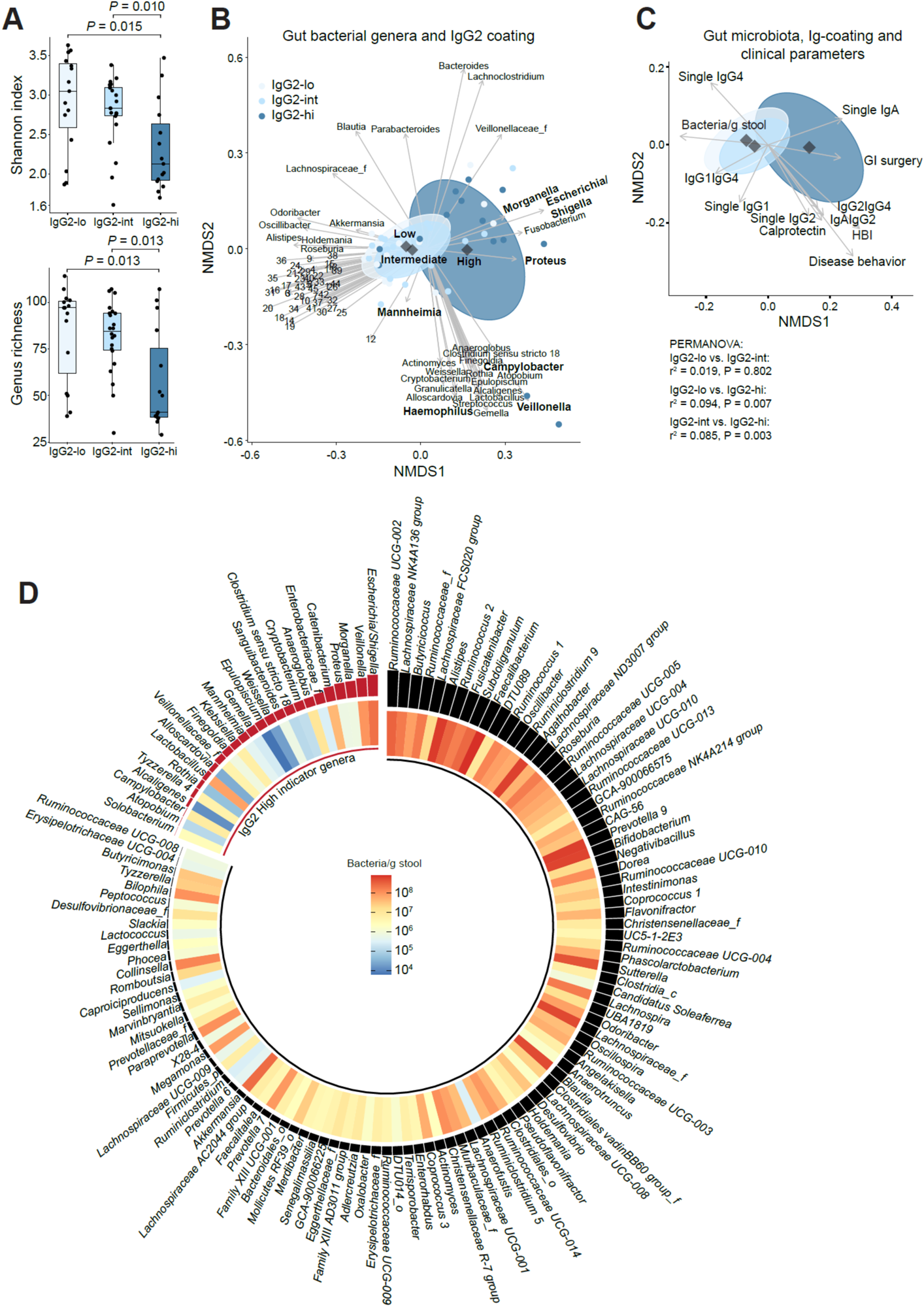
The gut microbiota of CD patients with high bacterial IgG2-coating associates with severe disease. a) Alpha-diversity of bacterial communities determined by Shannon index and genus richness within IgG2-coating subgroups. b) Non-metric multi-dimensional scaling (NMDS) plot, based on Bray-Curtis distances of the gut microbiota determined by 16S rRNA gene amplicon sequencing in CD patients. Individuals are represented as colored dots. Confidence ellipses represent the variance of the mean within each IgG2 subgroup. Individual gut bacterial taxa are fitted onto the plots and represented by arrows (see methods for details). Taxa names for the included numbers are provided in **Supplementary Table 3**. c) NMDS displaying Ig-coating levels and clinical parameters (instead of individual bacterial taxa). d) Heatmap showing the bacterial taxa and their abundance (number of bacteria/g stool) identified to separate the IgG2-hi subgroup from the IgG2-lo/int subgroups using sPLS-DA. Bars on top of each tile represent how important the individual bacterium is for the separation. Red bars highlight IgG2-hi indicator genera, while black bars highlight genera representing the IgG2-lo/int subgroups. For analysis, b-c) the length of arrows corresponds to r^2^-values (FDR-adjusted using q < 0.1) and each rhombus represent the center of the groups. Statistical analyses were based on Wilcoxon rank-sum test.

The composition of the overall gut microbiota in patients with IgG2-hi gut bacterial coating differed significantly from patients with IgG2-lo and IgG2-int as visualized using non-metric multidimensional scaling (NMDS) of the Bray-Curtis dissimilarity (**Figure 3b**, PERMANOVA: IgG2-hi vs IgG2-lo: r^2^ = 0.094, *P* = 0.007, IgG2-hi vs IgG2-int: r^2^ = 0.085, *P* = 0.003, **Supplementary Table 3**). Amongst the bacteria associated with IgG2-hi coating, we found several bacteria previously reported to be associated with CD, such as *Escherichia*/*Shigella, Veillonella, Morganella*, *Proteus*, *Campylobacter*, *Haemophilus,* and *Mannheimia*^17–19^. By correlating Ig-coating- and clinical parameters for CD patients with bacterial β-diversity patterns, we identified several bacterial species associated with IgG2-hi coating to follow a similar direction as several disease severity parameters (GI surgery, HBI and disease behavior) and enhanced IgA-coating, and to inversely relate to gut bacterial coating with single IgG1 and IgG4, and IgG1IgG4 double-coating (**Figure 3c**).

We confirmed by a Procrustes analysis that the distribution of all Ig-coating data and the distribution of the total bacterial community showed comparable patterns (Ig-coating (**Figure 1h**) vs total community β-diversity (**Figure 3b**, m1,2 = 0.386, *P* = 0.001). Since single- and double-IgG2-coating were main drivers of these patterns, and overlapped with disease severity parameters, these findings implied that IgG2-coating is the main driver of gut community patterns distinguishing severity of disease in CD patients.

We next performed a sparse partial least squared discriminant analysis (sPLS-DA)^20^ to identify bacterial indicator genera within the overall gut microbiota that are enriched in IgG2-hi vs IgG2-lo and IgG2-int. The sPLS-DA represents a cross-validated supervised clustering algorithm capable of identifying features important for separation of these groups. The model showed good predictive power (**Supplementary Figure 3b** [AUC: 0.827]) and resulted in the identification of 25 bacterial indicator genera that characterized the gut microbiota of IgG2-hi patients (**Figure 3d, Supplementary Table 4,** q < 0.1). The sPLS-DA-identified IgG2-hi indicator genera strongly overlapped with the disease severity-associated bacteria identified using the above NMDS, including *Escherichia*/*Shigella, Veillonella, Morganella*, *Proteus*, *Campylobacter*, *Haemophilus,* and *Mannheimia*.

### Identifying the nature of IgG2-coated gut bacteria in CD patients

FACS was next used to sort out IgG2-hi gut bacteria followed by sequencing of the bacterial V3-V4 16S rRNA gene region. This resulted in identification of 84 uniquely IgG2-coated genera out of 153 genera found in bulk stool from IgG2-hi CD patients (**Figure 4a** [inner circle vs. outer circle], **Supplementary Table 5 and 6**). When comparing the nature of the IgG2-coated bacteria with the identified IgG2-high indicator genera, it appeared that only 48% of the IgG2-hi indicator genera were IgG2-coated, meaning that some of the IgG2-hi indicator genera were non-coated. The non-coated bacteria in IgG2-hi CD patients included the Proteobacteria *Mannheimia*, *Morganella*, *Proteus*, *Campylobacter*, *Alcaligenes*, and members of the *Enterobacteriaceae*, while *Veillonella*, *Escherichia/Shigella*, *Klebsiella* and *Haemophilus* were IgG2-coated.

**Figure 4.**
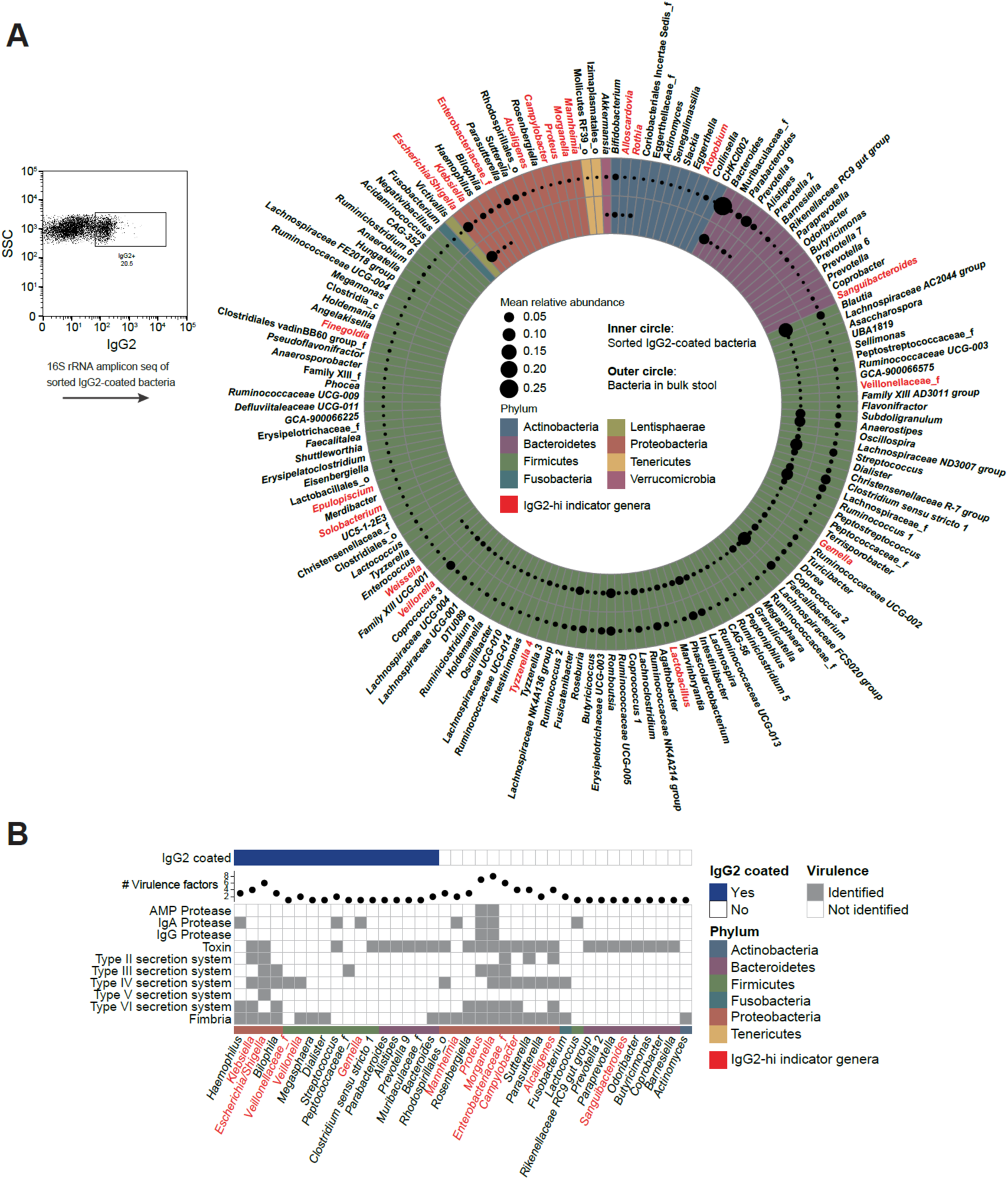
Identification of IgG2-coated gut bacteria in CD patients. a) IgG2-coated bacteria were sorted using FACS, and taxa were determined by 16S rRNA gene amplicon sequencing. Bacteria in bulk stool were sequenced from the same individuals, and the mean relative abundance of bacteria (dot size) in bulk stool (outer circle) and sorted IgG2-coated bacteria (inner circle) from CD patients within the IgG2-hi subgroup was determined. Data are shown at genus level with the phylum-level as tile color. b) *In silico* analysis of the presence of virulence factors important for invasion and immune evasion (grey square), in bacteria from a) with at least one virulence factor. Bacteria identified as IgG2-coated are marked with a blue square in the upper panel and the total number of virulence factors is presented as a dot in the middle panel. a-b) Bacteria highlighted in red represent IgG2-hi indicator genera identified in Figure 3.

To improve our understanding of functional differences and pathogenic potential between IgG2-hi coated and non-coated taxa, we performed *in silico* genome-based functional assessments using Picrust2^21^ for functional imputation based on taxonomy and the virulence factor database (VFDB)^22^ for identification of virulence factors important for invasion, immune evasion, and adherence (**Figure 4b**). Amongst the bacteria in bulk stool from CD patients, we identified 38 bacterial genera harboring at least one relevant virulence factor, amongst which 12 were IgG2-hi indicator genera, 17 were IgG2-coated, and 21 non-coated. It is notable that the non-coated Proteobacteria *Morganella*, *Proteus*, *Campylobacter*, and *Enterobacteriaceae* contained most virulence factors. We earlier identified these bacteria to be associated with severe disease and to represent IgG2-hi CD patients. Based on the presence of genes encoding all enzymes required for specific microbial biosynthetic pathways, we performed *in silico* prediction of the capability for flagella, hexa- and penta-acylated lipopolysaccharide (LPS) production in the bacteria, and found either flagella or hexa-acylated LPS to be present in the above non-coated bacteria identified in IgG2-hi CD patients (**Supplementary Figure 4**). Flagellin in flagella and hexa-acylated LPS are microbial ligands known for stimulating the immune system via toll-like receptor 5 (TLR5) and TLR4 activation, respectively^23, 24^, while penta-acylated LPS acts as a sequester of the TLR4 co-receptor myeloid differentiation factor 2 (MD-2), and diminishes human TLR4 activation^25, 26^.

### Distinctly co-occurring IgG2-coated and non-coated bacteria prevail in active disease

We next analyzed for presence of IgG2-hi gut bacteria with #virulence factors ≥1 in active vs. remissive disease status in the IBDSL cohort from The Netherlands (NL). *Campylobacter*, *Haemophilus*, *Mannheimia* and *Veillonella* were significantly enriched in individuals with active disease, while *Parasutterella*, *Lactococcus*, the *Muribaculaceae* family and the *Rhodospirillales* order were enriched in patients with remissive disease status (NL, **Figure 5a**). We performed a replication of this analysis in an independent cohort from the US (N = 297)^27^, and likewise found *Haemophilus*, *Campylobacter* and *Mannheimia* to associate with an active disease state (**Figure 5a**).

**Figure 5.**
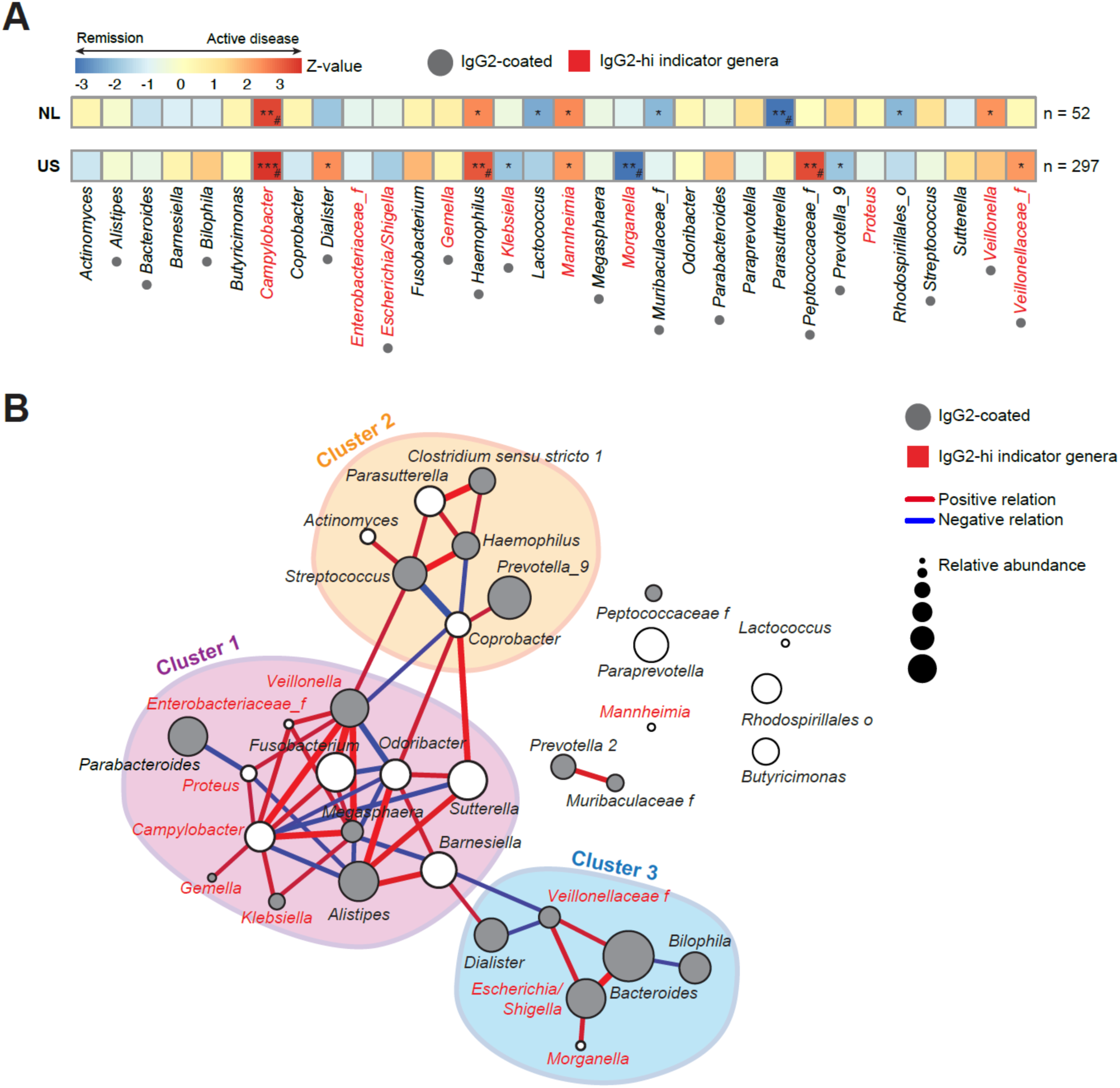
Distinct IgG2-coated gut bacteria and non-coated gut pathobionts co-exists in CD patients with active disease. a) Heatmap representing enrichment of identified IgG2-hi related bacteria harboring at least one virulence factor during active or remissive disease in CD patients from the IBDSL cohort from the Netherlands (NL) and an independent cohort from the United States (US). Z-values represent test statistics of coefficients for generalized linear models modelled over a negative binomial distribution. *P*-values are marked with stars: *, *P* < 0.05, **, *P* < 0.01, ***, *P* < 0.001, while # highlights taxa significant after FDR-adjustment, q < 0.05. b) Co-occurrence network of bacteria identified in CD patients from a) with active disease (n=19) in the NL cohort, where the taxonomy of IgG2-coated bacteria was determined. Clusters were identified using the walktrap algorithm and are indicated by an orange, blue or purple circle. Red and blue edges represent positive and negative relations, respectively. Node size is scaled by the relative abundance, and grey dots indicate that the bacterium is IgG2-coated. Bacteria highlighted in red represent IgG2-hi indicator genera identified in Figure 3.

Co-occurrence network analyses revealed distinct interconnections between the IgG2-hi gut bacteria with #virulence factors ≥1, as we discovered the existence of three different gut bacterial clusters in CD patients with active disease in the NL cohort (**Figure 5b**). Cluster 1 consisted of several known pathogens and genera implicated in CD, like *Klebsiella*, *Campylobacter*, *Proteus*, *Veillonella* and *Fusobacterium*^17–19, 28^, all identified as IgG2-hi indicator genera. Cluster 2 harbored several genera normally found in the oral cavity like *Streptococcus*, *Actinomyces*, *Haemophilus* and *Prevotella*^29^, none of which were identified to be IgG2-hi indicator genera. Cluster 3 included genera commonly found in the intestine, and among them were three IgG2-hi indicator genera, *Escherichia/Shigella*, *Morganella* and *Veillonellaceae*. All three clusters contained IgG2-coated bacteria, but in cluster 1, we also identified the active-disease associated and IgG2-hi indicator genera *Campylobacter*, which was found to be non-coated, to co-exist with several IgG2-coated bacteria. This co-existence between IgG2-coated and non-coated bacteria may explain why *Campylobacter* was identified as an IgG2-hi indicator genera, despite being non-coated. The analogous co-occurrence network analysis for the US cohort revealed a similar network structure of one cluster with mostly IgG2-hi indicator genera like *Klebsiella*, *Mannheimia, Proteus*, *Veillonella* and *Fusobacterium* (cluster 1), one cluster with genera often found in the oral community (cluster 2), and one cluster with bacteria often found in the gut (cluster 3) (**Supplementary Figure 5**). Interestingly, *Campylobacter* was part of the main network in CD patients with active disease from NL, but not in the US cohort, while the opposite was the case for *Mannheimia*, despite both being significantly associated to disease activity in both cohorts. These findings point to IgG2-hi indicator genera like the non-coated *Campylobacter* and *Mannheimia* as important, and apparently mutually exclusive drivers of active disease in CD patients, as we do not find them to exist together.

## Discussion

Understanding the natural dynamics of microbe-host interactions at mucosal surfaces might help in identifying new means of treatment for individuals with microbe-driven inflammatory diseases, including IBD. We here demonstrated that on average 1.7% of gut bacteria, corresponding to ca. 1.9×10^8^ bacteria/g stool, are coated with IgG in healthy individuals and CD patients. This fraction is only two to three-fold lower than the number of IgA-coated gut bacteria in healthy individuals and CD patients, indicating that IgG may hold a yet unrecognized role in regulating microbial dynamics in the gut. Although we found no overall significant differences in gut bacterial IgG-coating between healthy individuals and CD patients, we identified a subgroup of patients with severe CD and enhanced IgG2-coating that harbored a distinct microbiota with several gut pathobionts as indicator genera. Amplicon sequencing of the sorted IgG2-coated bacteria showed that IgG2-coating was quite promiscuous in nature, and coated 84 out of the 153 genera found in CD patients with IgG2-coating of gut bacteria. It is notable that we found less than 40% of IgG2-hi indicator genera to be IgG2-coated and that the IgG2-hi indicator genera with increased virulence potential were identified as non-coated. The two non-coated IgG2-hi indicator species *Campylobacter* and *Mannheimia* associated strongly with active disease in the NL cohort, which replicated in a US cohort. *Campylobacter* and *Mannheimia* were identified to co-cluster with bacteria in microbial cluster 1, in either the NL or the US cohort, where they co-existed with genera like *Veillonella* that represent a top-tier IgG2-coated IgG2-hi indicator genus. *Veillonella*, together with the other IgG2-hi indicator genera, *Klebsiella*, *Proteus* and *Escherichia*/*Shigella* have previously been associated with CD^29, 30, 40, 41^. We speculate that IgG2-coating of gut bacteria could be a means for the host to delimit specific bacterial growth and invasion in individuals harboring highly virulent bacteria, while the lack of IgG2-coating of certain gut pathobionts might be due to specific yet uncovered immune evasion mechanisms existing in these non-coated bacteria. The latter may leave a way for propagating pro-inflammatory responses, and thus disease flares in CD, as our findings imply. Targeted treatments with e.g. antibodies that bind to adhesion molecules on the two non-coated IgG2-hi indicator genera *Campylobacter* or *Mannheimia* to diminish their interaction with the intestinal epithelium may result in reduced mucosal invasion and inflammation, and thereby lessen disease activity. A few previous studies have reported that IgG’s are generated against gut bacteria by demonstrating the binding of serum-derived IgG’s to a panel of gut bacteria, and it has also been reported that IgG responses are generated against microbes in both healthy individuals^8^ and in patients suffering from autoimmune disorders and CD^30^. These previous studies demonstrated a profound overlap between the gut bacterial taxa that are recognized by serum-derived IgA and IgG, as we do in the present study by identification of double-coated bacteria. Moreover, it was earlier demonstrated that serum-derived IgG2 can bind to gut-derived bacteria, which was further supported by the presence of 35.9% of IgG2+ plasmablasts in terminal ileum^31^. These previous data support our findings that IgG2 can bind to both enteropathogenic and commensal bacteria in the human gut. One recent study that profiled the coating of gut bacteria with IgA, IgM, and IgG, identified a significant increase in the % of IgG2-coated bacteria in CD patients compared to healthy controls^32^. Although their study did not link findings to disease severity parameters nor identified the nature of IgG2-coated bacteria, our combined findings point to a role of IgG2-coating as a biomarker for CD patients with severe disease in which specific bacterial targeting with antibodies may be a way forward to relieve disease symptoms.

We identified two major non-coinciding gut pathobionts (*Mannheimia* and *Campylobacter*) as IgG2-hi indicator genera enriched in patients with active CD that remained non-coated despite high IgG2-coating of other co-existing bacteria, like *Veillonella*. *Mannheimia* has mostly been described as an animal pathogen infecting the airways, but cases of humans being infected by *Mannheimia* have been reported^33^. Our *in silico* functional analysis of *Mannheimia*’s virulence potential showed that it may have the capacity to produce IgA-specific proteases, hence escaping IgA coating. *Campylobacter* is a well-known human pathogen and is often the causative agent of food poisoning^34^, but has also been identified in the gut of CD patients^35, 36^. Because of the enteroinvasive nature and complex outer membrane of *Campylobacter*, they are often found to infect intestinal epithelial cells as a mean to evade humoral responses^34, 37^.

The difference between bacteria that are capable of inducing IgG2 responses and become IgG2-coated, and those that are able to induce IgG2 responses while avoiding Ig-coating might be a defining feature in disease-driving species, since IgG2 reactions are otherwise known for their effectiveness in clearing invasive pathogens, e.g. in patients with aggressive periodontitis^6^. It may be that invasive and/or toxin-producing bacteria, like *Campylobacter*, are able to initiate the breakdown of the intestinal integrity. This would increase influx of luminal antigens and co-existing gut bacteria into the underlying tissue and initiate a Type 1-immune reaction dominated by IL-12p70 and IFN-ɣ (due to immunostimulating ligands in *Campylobacter*), and thus subsequently, promote IgG2-production^17, 38, 39^. When the barrier is impaired, bacteria like *Escherichia*^40^ or *Klebsiella*^41^, both expressing the Type 1-immune activating ligand hexa-acylated LPS, may get in contact with immune cells in the lamina propria and prime a Type 1 immune response, thereby stimulating processes that may result in antigen-specific IgG2 production against *Escherichia* or *Klebsiella*, as well as other co-existing microorganisms.

Combined, we here demonstrated that gut bacterial IgG2-coating characterizes individuals with severe CD, and is enhanced during disease flares. In this patient group we identified the distinct presence of two non-coated gut pathobionts, *Campylobacter* or *Mannheimia*, that we speculate may drive the inflammatory processes, and thus enhance disease severity, as they may linger in an uncontrolled manner when non-coated. Specific therapeutic elimination of *Campylobacter* or *Mannheimia* may therefore be a strategy to relieve disease symptoms in IgG2-hi CD individuals with severe disease burden. The current findings point to new ways for subgrouping of CD patients by utilizing the host immune system in identifying disease-propagating bacteria and in pinpointing the underlying immune reactions directed against these bacteria.

## Acknowledgements

The authors thank the participants of the IBDSL and the Maastricht IBS cohorts, Maastricht, The Netherlands, for providing samples.

## Author contributions

Conceptual design of the study and method implementation: C.E., S.B., Responsible for the NL cohort and fecal 16S rRNA analysis: J.P., D.M.A.E.J. Generation of Ig-coating profiles: P.W.B., S.V., T.B.H., C.E. Data analysis: C.E. supported by J.M.M., P.N.M., S.B. Statistical support: P.R. Sorting of coated bacteria: C.E., L.B.R. supported by S.B. Implementation of volume-based method and sequencing of IgG2-coated bacteria: C.E. supported by N.B.D.S., K.K., S.B. Writing of draft manuscript: C.E., S.B. Critical revision of manuscript: all authors.

## Data Availability

Sequencing data are publicly available in NCBI Sequence Read Archive under BioProject: PRJNA418765. Analysis software including quality control, taxonomic, and functional inference tools are publicly available and referenced as appropriate.

## Abbreviations

CD: Crohn’s Disease
FACS: fluorescence-activated cell sorting
FDR: false discovery rate
HBI: Harvey-Bradshaw Index
IBD: Inflammatory bowel disease
IBDSL: IBD South Limburg
Ig: Immunoglobulin
LPS: lipopolysaccharide
NMDS: non-metric multidimensional scaling
sPLS-DA: sparse partial least squared discriminant analysis
TLR: toll-like receptor.

## Online Methods

### Cohort characteristics

Samples from CD patients (n=60) were collected as part of the IBD South Limburg (IBDSL) cohort, which is a population-based inception cohort from the South Limburg area of the Netherlands^12^. Since 1991, all newly diagnosed patients with IBD have been prospectively included and followed, with on-going collection of biomaterials (serum, plasma, DNA and stool), and complete data on disease phenotype, hospitalizations, surgery, (extra)intestinal complications, and diagnostic reports. We included CD patients who have not received antibiotics treatment two months before sampling. Samples were taken either during remission (n=33) or during active disease (n=27). Active disease was defined by clinicians as fecal calprotectin >250 μg/g or fecal calprotectin >100 μg/g and at least a five-fold increase from baseline. The cohort was sampled as described in the IBDSL cohort profile^12^. The control samples from healthy individuals (n=20) were collected as part of the Maastricht IBS cohort. Both the IBDSL and the Maastricht IBS cohort were approved by the local Medical Ethics Committee, registered in http://www.clinicaltrials.gov (NCT02130349 and NCT00775060), and follow the revised version of the declaration of Helsinki. The data used for replication derive from a previous publication from a US based cohort^27^. Briefly, raw data from treatment-naïve CD patients (i.e. no treatment with Ustekinumab) were processed using the DADA2 pipeline, and samples with >5,000 reads were used in the final analysis, resulting in inclusion of 297 samples. Fecal calprotectin >250 μg/g was defined as active disease.

### Determination of fecal bacterial load and bacterial Ig-coating by flow cytometry

Fecal samples were incubated on ice for 1 hour in sterile PBS at 100 mg/mL, homogenized, spun down (15 min, 50g, 4°C), followed by aspiration of the supernatant. The supernatant was centrifuged (5 min, 8000g, 4°C) and washed twice in buffer 1 (PBS + 1% BSA (>98%, Sigma-Aldrich)). An aliquot was diluted 150x in buffer 2 (PBS + 1% BSA + 0.01% Tween 20 + 1 mM EDTA) with 1 mM DAPI and 10 µL count beads (BD Biosciences) and analyzed on a FACS Canto II flow cytometer (BD Biosciences). The bacterial population was gated based on SSC-A/Pacific-Blue (**Supplementary Figure 1**), and the fecal bacterial load was determined as per instruction by the bead manufacturer. For consistent staining across samples, a total of 1.5×10^8^ bacteria per sample were transferred to a new tube. The bacteria were resuspended in buffer 1 containing 20% mouse serum and incubated for 20 min at 4°C. The percentage of bacterial coating with IgA and IgG1-4 was defined by staining of samples for 30 min at 4°C with a mixture of fluorescence-conjugated mouse anti-human Ig antibodies that each bind to the constant part of the human antibodies: Pe-Cy7-anti-IgA (Miltenyi), Biotin-anti-IgG1 + APC-Cy7-Streptavidin (SouthernBiotech), Alexa Flour 647-anti-IgG2 (SouthernBiotech), Alexa Flour 488-anti-IgG3 (SouthernBiotech), and PE-anti-IgG4 (SouthernBiotech) diluted in buffer 1. Samples were washed twice in buffer 1 and resuspended in buffer 2. An aliquot was diluted and incubated with 1 mM DAPI and 10 µL count beads (BD Biosciences) before being analyzed on a FACS Canto II flow cytometer (BD Biosciences). The analysis was based on 200,000 recorded DAPI^+^ cells. Data were analyzed using FlowJo software (Version 10.5.0, Tree Star Inc, Ashland, OR). All gate boundaries were set using FMO controls.

### Purification of IgG2-coated bacteria by FACS

Samples for sorting of IgG2-coated bacteria were prepared as described above, except for the use of PE-conjugated mouse anti-human IgG2 (0.5 mg/mL, SouthernBiotech). Samples were sorted on a MoFlo XDP Cell sorter (Beckman Coulter). The IgG2+ fraction (between 1.5×10^5^ to 3×10^6^ bacteria per sample) was collected in heat-inactivated fetal bovine serum (FBS; Gibco) coated FACS tubes, pelleted (5 min, 8000g, 4°C) and stored at -80°C until processing. Samples of sheath fluid were collected directly from the stream pre-sorting as technical controls.

### DNA extraction and 16S rDNA library preparation of IgG2-coated bacteria

Bacterial DNA was extracted using the NucleoSpin Soil kit (Macherey-Nagel, Germany) based on the manufacturer’s protocol. The extracted DNA was amplified using a two-step PCR reaction with the 314F (TCGTCGGCAGCGTCAGATGTGTATAAGAGACAGCCTACGGGNGGCWGCAG) and 806R (GTCTCGTGGGCTCGGAGATGTGTATAAGAGACAGGACTACHVGGGTATCTAATCC) targeting the hypervariable regions V3 + V4 of the 16S ribosomal RNA gene. DNA was amplified using Phusion Green High-Fidelity DNA Polymerase (Thermo Fisher) kit. For PCR, master mix was added to 20 µL extracted DNA in concentrations according to manufacturer’s recommendations. PCR was performed using the following conditions: initial denaturation for 30s at 98°C followed by 28 cycles of 10s 98°C, 15s 56°C and 30s 72°C with a final elongation at 72°C for 5 min. The products were tagged with Illumina adapters (Forward: AATGATACGGCGACCACCGAGATCTACAC, Reverse: CAAGCAGAAGACGGCATACGAGAT) using 10 cycles under the same PCR conditions. Products from both PCRs were purified using Agencourt AMPure XP beads (Beckman Coulter). Similar volumes of all amplicons were pooled and the library was diluted to a total concentration of 4 nM before sequencing on the MiSeq platform (Illumina, USA) using the v3 kit (paired-end).

### 16S rRNA gene data processing

Sequencing adapters were removed using the BBDuk tool in the BBTools package v38.37 (BBDuk, sourceforge.net/projects/bbmap/). Reads were analyzed and denoised using DADA2^42^. Resulting amplicon sequence variants (ASVs) were compared to the 99% identity clustered SILVA database v132^43^ using a naive bayesian classifier^44^ trained on the amplified region as implemented in DADA2. Since samples and technical controls were adjusted to sample-volume rather than DNA concentration in the PCR, the false-positive count contribution originating from the technical controls, is similar between samples. This enabled us to correct for reagent and pre-sorting fluid contaminating bacterial DNA by subtracting the read counts found in the above technical controls. Bacterial reads data are found in **Supplementary Table 5.**

### *In silico*-based inference of virulence factors and immunostimulatory ligands production capability in gut bacteria

Bacteria were annotated for virulence factors and capacity to produce hexa- or penta-acylated LPS, and flagellin by predicting their functional capacity using PICRUST2^21^ (v. 2.4.1) to identify Kyoto Encyclopedia of Genes and Genomes (KEGG) orthologs (KOs)^45^. For each ASV, flagellin production was evaluated by the presence of K numbers responsible for production of flagellin as described in the flagellar assembly map (map02040). The type of LPS, or whether a bacterium could produce it at all, was evaluated by the ability of each species to convert UDP-N-acetylglucosamine to KDO_2_-lipid A as described in the lipopolysaccharide biosynthesis pathway (module M00060)^23^. For virulence factors, we downloaded protein sequences from the Virulence Factors Database (VFDB)^22^ and mapped the sequences to K numbers using GhostKOALA (https://www.kegg.jp/ghostkoala)^45^. Resulting KOs were integrated with the predicted functions from the PICRUST2 analysis to assign information on the virulence potential to each bacterium in our samples.

## Statistical analysis

Statistical analyses were performed using R (v. 4.0.0). The Shannon index and species richness were calculated using the vegan package (v. 2.6-2). Bray-Curtis dissimilarity measures were used to compute differences in bacterial communities within the IgG2 subgroups and presented using an NMDS based on seven dimensions. For the latter, the metaMDS function from the vegan package was used to iteratively add dimensions until lowest level of stress was achieved; taxa or clinical variables and Ig-coating were added to the NMDS using the envfit function. Group differences were tested for inference using a permutational multivariate analysis of variance (PERMANOVA with 999 permutations, using the Adonis2 function from the vegan package). To identify high IgG2 indicator species within bulk fecal sequences, we trained models of sPLS-DA using the mixomics package (v. 6.2)^20^ on the log10-transformed relative abundances, using 1/2 of the lowest non-zero value as pseudocounts. The optimal number of species was identified by 10-fold cross-validation using AUC statistics to avoid overfitting the models. Disease activity was fitted to bacterial counts from virulent genera (defined as number of virulence factors ≥ 1) using negative binomial generalized linear models with logged sequencing depths as offset-term and using the NBZIMM package (v. 1.0)^46^. Microbiota community structures in CD patients with active disease were evaluated by building co-occurrence networks of virulent species (#virulence factors ≥ 1) using Sparse Correlations for Compositional data (SparCC) algorithm^47^ from the SpiecEasi package (v. 1.1.2)^48^. Bacteria-bacteria correlation coefficients were estimated as the average of 100 inference iterations refined by 999 exclusion iterations with a strength threshold of 0.6. Correlation coefficients with an absolute value of 0.3 or above were visualized. Clustering was done using the walktrap community algorithm from the igraph package (v. 1.3.2) using default parameters.

Statistical non-parametric tests were used for all data comparisons: the Wilcoxon rank-sum test was used when comparing two groups, and Spearman rank coefficient correlations were used for association analysis. *P*-values were deemed significant using *P* < 0.05 as significance level. When indicated, *P*-values were adjusted for multiple testing using a false discovery rate (FDR) adjustment of q < 0.10 or < 0.05.

