## Supplementary Tables for "Specific gut pathobionts escape antibody coating and are enriched during flares in patients with severe Crohn’s disease"

**Supplementary Table 1 - Cohort statistics**

| | Healthy | CD patients |
|---|---|---|
| **Basic characteristics** | | |
| Sex, Female, % (N) | 65% (13) | 66% (40) |
| Age, mean (SD) | 41 (17) | 42 (15) |
| **Clinical parameters** | | |
| Disease activity, Active, % (N) | - | 45% (27) |
| Fecal calprotectin (μg/g), mean (SD) | - | 220.68 (307.25) |
| HBI, mean (SD) | - | 2.92 (3.41) |
| Surgery, % (Total N) | - | 20% (58) |
| none, N | - | 46 |
| ileocoecaal, N | - | 4 |
| colon, N | - | 3 |
| hemicolectomy, N | - | 3 |
| sigmoid, N | - | 1 |
| rectum+sigmoid, N | - | 1 |
| **Montreal Classification** | - | |
| Disease Location (Total N) | - | 60 |
| L1, % (N) | - | 32% (19) |
| L2, % (N) | - | 28% (17) |
| L3, % (N) | - | 40% (24) |
| Disease behavior (Total N) | - | 60 |
| B1, % (N) | - | 66.7% (40) |
| B2, % (N) | - | 20% (12) |
| B3, % (N) | - | 13% (8) |
| Age at diagnosis (Total N) | - | 60 |
| A1, % (N) | - | 3% (2) |
| A2, % (N) | - | 75% (45) |
| A3, % (N) | - | 22% (13) |

### Supplementary Table 2 - Quantitative coating data

**Ig-coated bacteria per g stool in healthy controls vs. CD (Wilcoxon Rank Sum test)**

| Isotype | Donor type | Data type | Median | 25th quantile | 75th quantile | Donor type | Data type | Median | 25th quantile | 75th quantile | P-value | Adjusted (FDR) | Significance | Method |
|---|---|---|---|---|---|---|---|---|---|---|---|---|---|---|
| Single IgA | Control | Bacteria per g stool | 5.89E+08 | 2.05E+08 | 8.60E+08 | CD | Bacteria per g stool | 6.45E+08 | 4.20E+08 | 1.34E+09 | 1.73E-01 | 8.00E-01 | ns | Wilcoxon |
| Single IgG1 | Control | Bacteria per g stool | 5.65E+07 | 3.53E+07 | 8.96E+07 | CD | Bacteria per g stool | 3.85E+07 | 1.78E+07 | 1.01E+08 | 2.28E-01 | 8.00E-01 | ns | Wilcoxon |
| Single IgG2 | Control | Bacteria per g stool | 0 | 0 | 7.25E+05 | CD | Bacteria per g stool | 2.07E+04 | 0 | 1.67E+06 | 2.15E-01 | 8.00E-01 | ns | Wilcoxon |
| Single IgG3 | Control | Bacteria per g stool | 6.59E+05 | 0 | 1.77E+06 | CD | Bacteria per g stool | 0 | 0 | 9.82E+05 | 9.68E-02 | 8.00E-01 | ns | Wilcoxon |
| Single IgG4 | Control | Bacteria per g stool | 1.03E+08 | 6.41E+07 | 2.57E+08 | CD | Bacteria per g stool | 8.02E+07 | 2.64E+07 | 2.74E+08 | 5.67E-01 | 8.40E-01 | ns | Wilcoxon |
| IgAIgG1 | Control | Bacteria per g stool | 1.72E+07 | 5.07E+06 | 2.43E+07 | CD | Bacteria per g stool | 1.17E+07 | 5.53E+06 | 2.66E+07 | 6.85E-01 | 9.10E-01 | ns | Wilcoxon |
| IgAIgG2 | Control | Bacteria per g stool | 1.10E+06 | 0 | 3.05E+06 | CD | Bacteria per g stool | 6.65E+05 | 0 | 5.78E+06 | 9.45E-01 | 1.00E+00 | ns | Wilcoxon |
| IgAIgG3 | Control | Bacteria per g stool | 0 | 0 | 9.69E+05 | CD | Bacteria per g stool | 0 | 0 | 0.00 | 6.51E-02 | 8.00E-01 | ns | Wilcoxon |
| IgAIgG4 | Control | Bacteria per g stool | 8.15E+06 | 5.26E+06 | 1.72E+07 | CD | Bacteria per g stool | 9.17E+06 | 4.30E+06 | 1.94E+07 | 9.78E-01 | 1.00E+00 | ns | Wilcoxon |
| IgG1IgG2 | Control | Bacteria per g stool | 0 | 0 | 3.63E+05 | CD | Bacteria per g stool | 0 | 0 | 0.00 | 4.61E-01 | 8.40E-01 | ns | Wilcoxon |
| IgG1IgG3 | Control | Bacteria per g stool | 0 | 0 | 0 | CD | Bacteria per g stool | 0 | 0 | 0.00 | 2.62E-01 | 8.10E-01 | ns | Wilcoxon |
| IgG1IgG4 | Control | Bacteria per g stool | 1.54E+06 | 7.80E+05 | 2.29E+06 | CD | Bacteria per g stool | 1.14E+06 | 3.92E+05 | 2.92E+06 | 4.80E-01 | 8.40E-01 | ns | Wilcoxon |
| IgG2IgG3 | Control | Bacteria per g stool | 0 | 0 | 0 | CD | Bacteria per g stool | 0 | 0 | 0.00 | 9.77E-01 | 1.00E+00 | ns | Wilcoxon |
| IgG2IgG4 | Control | Bacteria per g stool | 0 | 0 | 7.10E+04 | CD | Bacteria per g stool | 0 | 0 | 0.00 | 7.94E-01 | 9.70E-01 | ns | Wilcoxon |
| IgG3IgG4 | Control | Bacteria per g stool | 0 | 0 | 5.97E+05 | CD | Bacteria per g stool | 0 | 0 | 3.00E+05 | 5.31E-01 | 8.40E-01 | ns | Wilcoxon |
| IgAIgG1IgG2 | Control | Bacteria per g stool | 0 | 0 | 5.13E+05 | CD | Bacteria per g stool | 0 | 0 | 1.45E+05 | 7.61E-01 | 9.70E-01 | ns | Wilcoxon |
| IgAIgG1IgG4 | Control | Bacteria per g stool | 5.24E+05 | 0 | 1.31E+06 | CD | Bacteria per g stool | 3.42E+05 | 0 | 1.07E+06 | 5.92E-01 | 8.40E-01 | ns | Wilcoxon |
| IgAIgG2IgG3 | Control | Bacteria per g stool | 0 | 0 | 0 | CD | Bacteria per g stool | 0 | 0 | 0 | 5.83E-01 | 8.40E-01 | ns | Wilcoxon |
| IgAIgG2IgG4 | Control | Bacteria per g stool | 0 | 0 | 0 | CD | Bacteria per g stool | 0 | 0 | 0 | 1.00E+00 | 1.00E+00 | ns | Wilcoxon |
| IgAIgG3IgG4 | Control | Bacteria per g stool | 0 | 0 | 0 | CD | Bacteria per g stool | 0 | 0 | 0 | 1.39E-01 | 8.00E-01 | ns | Wilcoxon |
| IgG1IgG2IgG4 | Control | Bacteria per g stool | 0 | 0 | 0 | CD | Bacteria per g stool | 0 | 0 | 0 | 4.23E-01 | 8.40E-01 | ns | Wilcoxon |
| IgG1IgG3IgG4 | Control | Bacteria per g stool | 0 | 0 | 0 | CD | Bacteria per g stool | 0 | 0 | 0 | 4.35E-01 | 8.40E-01 | ns | Wilcoxon |
| IgAIgG1IgG2IgG4 | Control | Bacteria per g stool | 0 | 0 | 0 | CD | Bacteria per g stool | 0 | 0 | 0 | 3.26E-01 | 8.40E-01 | ns | Wilcoxon |

#### Supplementary Table 3 - Taxa annotation for numbers on NMDS plot

| Genus | NMDS1 | NMDS2 | r | pvals | pvals_adj | Numbering on plot |
|---|---|---|---|---|---|---|
| *Clostridiales_o* | -0.353 | -0.117 | 0.283 | 6.00E-03 | 1.73E-02 | 1 |
| *Ruminococcaceae_UCG-003* | -0.449 | -0.066 | 0.420 | 1.00E-03 | 3.98E-03 | 2 |
| *GCA-900066575* | -0.492 | -0.139 | 0.533 | 1.00E-03 | 3.98E-03 | 3 |
| *Family_XIII_AD3011_group* | -0.403 | -0.064 | 0.340 | 1.00E-03 | 3.98E-03 | 4 |
| *Christensenellaceae_f* | -0.419 | -0.111 | 0.383 | 1.00E-03 | 3.98E-03 | 5 |
| *Subdoligranulum* | -0.494 | -0.138 | 0.536 | 1.00E-03 | 3.98E-03 | 6 |
| *Ruminococcaceae_UCG-010* | -0.393 | -0.138 | 0.355 | 1.00E-03 | 3.98E-03 | 7 |
| *Oscillospira* | -0.414 | -0.109 | 0.374 | 1.00E-03 | 3.98E-03 | 8 |
| *Lachnospiraceae_ND3007_group* | -0.415 | -0.031 | 0.354 | 1.00E-03 | 3.98E-03 | 9 |
| *Christensenellaceae_R-7_group* | -0.432 | -0.161 | 0.434 | 1.00E-03 | 3.98E-03 | 10 |
| *GCA-900066225* | -0.345 | -0.063 | 0.251 | 5.00E-03 | 1.51E-02 | 11 |
| *Bifidobacterium* | -0.198 | -0.282 | 0.243 | 1.00E-03 | 3.98E-03 | 12 |
| *Ruminococcaceae_UCG-009* | -0.334 | -0.056 | 0.234 | 1.00E-02 | 2.62E-02 | 13 |
| *Ruminococcus_1* | -0.487 | -0.225 | 0.588 | 1.00E-03 | 3.98E-03 | 14 |
| *Pseudoflavonifractor* | -0.344 | -0.046 | 0.246 | 3.00E-03 | 1.03E-02 | 15 |
| *Ruminococcaceae_UCG-002* | -0.539 | -0.129 | 0.627 | 1.00E-03 | 3.98E-03 | 16 |
| *Clostridiales_vadinBB60_group_f* | -0.498 | -0.115 | 0.533 | 1.00E-03 | 3.98E-03 | 17 |
| *Faecalibacterium* | -0.523 | -0.216 | 0.652 | 1.00E-03 | 3.98E-03 | 18 |
| *Lachnospiraceae_FCS020_group* | -0.486 | -0.242 | 0.602 | 1.00E-03 | 3.98E-03 | 19 |
| *Ruminococcaceae_f* | -0.563 | -0.185 | 0.717 | 1.00E-03 | 3.98E-03 | 20 |
| *Ruminiclostridium_5* | -0.478 | -0.076 | 0.478 | 1.00E-03 | 3.98E-03 | 21 |
| *CAG-56* | -0.385 | -0.084 | 0.316 | 1.00E-03 | 3.98E-03 | 22 |
| *Angelakisella* | -0.450 | -0.094 | 0.431 | 1.00E-03 | 3.98E-03 | 23 |
| *Ruminococcaceae_UCG-013* | -0.466 | -0.052 | 0.448 | 1.00E-03 | 3.98E-03 | 24 |
| *Lachnospira* | -0.300 | -0.199 | 0.265 | 2.00E-03 | 7.65E-03 | 25 |
| *Marvinbryantia* | -0.333 | -0.119 | 0.255 | 5.00E-03 | 1.51E-02 | 26 |
| *Agathobacter* | -0.358 | -0.167 | 0.319 | 1.00E-03 | 3.98E-03 | 27 |
| *Ruminococcaceae_NK4A214_group* | -0.451 | -0.148 | 0.460 | 1.00E-03 | 3.98E-03 | 28 |
| *Clostridia_c* | -0.430 | -0.072 | 0.388 | 1.00E-03 | 3.98E-03 | 29 |
| *Coprococcus_1* | -0.372 | -0.198 | 0.362 | 1.00E-03 | 3.98E-03 | 30 |
| *Ruminococcaceae_UCG-005* | -0.555 | -0.136 | 0.667 | 1.00E-03 | 3.98E-03 | 31 |
| *Adlercreutzia* | -0.334 | -0.160 | 0.280 | 1.00E-03 | 3.98E-03 | 32 |
| *Fusicatenibacter* | -0.381 | -0.104 | 0.318 | 1.00E-03 | 3.98E-03 | 33 |
| *Ruminococcus_2* | -0.471 | -0.188 | 0.524 | 1.00E-03 | 3.98E-03 | 34 |
| *Lachnospiraceae_NK4A136_group* | -0.546 | -0.091 | 0.626 | 1.00E-03 | 3.98E-03 | 35 |
| *Intestinimonas* | -0.515 | -0.036 | 0.544 | 1.00E-03 | 3.98E-03 | 36 |
| *Ruminococcaceae_UCG-014* | -0.386 | -0.167 | 0.362 | 1.00E-03 | 3.98E-03 | 37 |
| *Ruminiclostridium_6* | -0.334 | -0.020 | 0.228 | 9.00E-03 | 2.42E-02 | 38 |
| *Lachnospiraceae_UCG-008* | -0.318 | -0.068 | 0.216 | 5.00E-03 | 1.51E-02 | 39 |
| *Ruminiclostridium_9* | -0.440 | -0.092 | 0.412 | 1.00E-03 | 3.98E-03 | 40 |
| *DTU089* | -0.377 | -0.167 | 0.347 | 1.00E-03 | 3.98E-03 | 41 |
| *Lachnospiraceae_UCG-001* | -0.372 | -0.157 | 0.333 | 1.00E-03 | 3.98E-03 | 42 |
| *Negativibacillus* | -0.433 | -0.095 | 0.401 | 1.00E-03 | 3.98E-03 | 43 |
| *Coprococcus_3* | -0.323 | -0.109 | 0.237 | 3.00E-03 | 1.03E-02 | 44 |
| *Family_XIII_UCG-001* | -0.402 | -0.124 | 0.362 | 1.00E-03 | 3.98E-03 | 45 |

**Supplementary Table 4**

**Bacteria enriched in the IgG2 Phenotype as selected by sPLS-DA**

| **Genus** | **Importance** | **Frequency of festure selection in 5-fold cross-validation** |
|---|---|---|
| *Escherichia.Shigella* | 0.119 | 1.00 |
| *Veillonella* | 0.104 | 1.00 |
| *Morganella* | 0.092 | 0.99 |
| *Proteus* | 0.090 | 0.97 |
| *Catenibacterium* | 0.081 | 0.92 |
| *Enterobacteriaceae_f* | 0.060 | 0.91 |
| *Anaeroglobus* | 0.052 | 0.89 |
| *Clostridium_sensu_stricto_18* | 0.050 | 0.80 |
| *Cryptobacterium* | 0.050 | 0.80 |
| *Epulopiscium* | 0.050 | 0.80 |
| *Weissella* | 0.050 | 0.80 |
| *Sanguibacteroides* | 0.050 | 0.80 |
| *Gemella* | 0.047 | 0.88 |
| *Mannheimia* | 0.044 | 0.79 |
| *Klebsiella* | 0.039 | 0.81 |
| *Finegoldia* | 0.039 | 0.83 |
| *Veillonellaceae_f* | 0.036 | 0.84 |
| *Alloscardovia* | 0.026 | 0.82 |
| *Lactobacillus* | 0.025 | 0.74 |
| *Rothia* | 0.021 | 0.82 |
| *Tyzzerella_4* | 0.020 | 0.68 |
| *Alcaligenes* | 0.015 | 0.84 |
| *Campylobacter* | 0.002 | 0.76 |
| *Atopobium* | 0.001 | 0.64 |
| *Solobacterium* | 0.000 | 0.50 |
| *Ruminococcaceae_UCG.008* | -0.001 | 0.56 |
| *Erysipelotrichaceae_UCG.004* | -0.002 | 0.57 |
| *Butyricimonas* | -0.003 | 0.50 |
| *Tyzzerella* | -0.003 | 0.46 |
| *Bilophila* | -0.004 | 0.54 |
| *Peptococcus* | -0.004 | 0.53 |
| *Desulfovibrionaceae_f* | -0.004 | 0.43 |
| *Slackia* | -0.006 | 0.50 |
| *Lactococcus* | -0.008 | 0.55 |
| *Eggerthella* | -0.008 | 0.57 |
| *Phocea* | -0.010 | 0.63 |
| *Collinsella* | -0.010 | 0.62 |
| *Romboutsia* | -0.011 | 0.50 |
| *Caproiciproducens* | -0.013 | 0.78 |
| *Sellimonas* | -0.013 | 0.61 |
| *Marvinbryantia* | -0.014 | 0.62 |
| *Mitsuokella* | -0.015 | 0.90 |
| *Prevotellaceae_f* | -0.016 | 0.86 |
| *Paraprevotella* | -0.016 | 0.68 |

| | | |
|---|---|---|
| *X28.4* | -0.017 | 0.83 |
| *Megamonas* | -0.017 | 0.78 |
| *Lachnospiraceae_UCG.009* | -0.018 | 0.82 |
| *Firmicutes_p* | -0.020 | 0.76 |
| *Ruminiclostridium* | -0.024 | 0.95 |
| *Prevotella_6* | -0.024 | 0.96 |
| *Akkermansia* | -0.024 | 0.76 |
| *Lachnospiraceae_AC2044_group* | -0.028 | 0.91 |
| *Faecalitalea* | -0.029 | 0.80 |
| *Prevotella_7* | -0.031 | 0.92 |
| *Family_XIII_UCG.001* | -0.031 | 0.82 |
| *Bacteroidales_o* | -0.031 | 0.96 |
| *Mollicutes_RF39_o* | -0.034 | 0.96 |
| *Merdibacter* | -0.036 | 0.87 |
| *Senegalimassilia* | -0.038 | 0.93 |
| *GCA.900066225* | -0.039 | 0.90 |
| *Eggerthellaceae_f* | -0.040 | 0.88 |
| *Family_XIII_AD3011_group* | -0.040 | 0.91 |
| *Adlercreutzia* | -0.042 | 0.92 |
| *Oxalobacter* | -0.043 | 0.99 |
| *Erysipelotrichaceae_f* | -0.044 | 0.92 |
| *Ruminococcaceae_UCG.009* | -0.048 | 0.96 |
| *DTU014_o* | -0.049 | 1.00 |
| *Terrisporobacter* | -0.049 | 0.95 |
| *Enterorhabdus* | -0.051 | 1.00 |
| *Coprococcus_3* | -0.051 | 0.92 |
| *Actinomyces* | -0.053 | 0.95 |
| *Christensenellaceae_R.7_group* | -0.057 | 0.96 |
| *Muribaculaceae_f* | -0.058 | 1.00 |
| *Lachnospiraceae_UCG.001* | -0.063 | 0.98 |
| *Anaerofustis* | -0.065 | 1.00 |
| *Ruminiclostridium_5* | -0.066 | 1.00 |
| *Ruminococcaceae_UCG.014* | -0.066 | 1.00 |
| *Clostridiales_o* | -0.068 | 1.00 |
| *Pseudoflavonifractor* | -0.071 | 1.00 |
| *Holdemania* | -0.073 | 1.00 |
| *Desulfovibrio* | -0.073 | 1.00 |
| *Lachnospiraceae_UCG.008* | -0.076 | 1.00 |
| *Clostridiales_vadinBB60_group_f* | -0.080 | 1.00 |
| *Blautia* | -0.080 | 0.97 |
| *Anaerotruncus* | -0.081 | 1.00 |
| *Angelakisella* | -0.081 | 1.00 |
| *Ruminococcaceae_UCG.003* | -0.086 | 1.00 |
| *Oscillospira* | -0.088 | 1.00 |
| *Lachnospiraceae_f* | -0.088 | 0.97 |
| *Odoribacter* | -0.089 | 1.00 |
| *UBA1819* | -0.090 | 1.00 |
| *Lachnospira* | -0.090 | 0.98 |

| | | |
|---|---|---|
| *Candidatus_Soleaferrea* | -0.091 | 1.00 |
| *Clostridia_c* | -0.095 | 1.00 |
| *Sutterella* | -0.095 | 1.00 |
| *Phascolarctobacterium* | -0.096 | 1.00 |
| *Ruminococcaceae_UCG.004* | -0.097 | 1.00 |
| *UC5.1.2E3* | -0.098 | 1.00 |
| *Christensenellaceae_f* | -0.099 | 1.00 |
| *Flavonifractor* | -0.106 | 1.00 |
| *Coprococcus_1* | -0.106 | 1.00 |
| *Intestinimonas* | -0.108 | 1.00 |
| *Ruminococcaceae_UCG.010* | -0.109 | 1.00 |
| *Dorea* | -0.111 | 0.99 |
| *Negativibacillus* | -0.119 | 1.00 |
| *Bifidobacterium* | -0.124 | 1.00 |
| *Prevotella_9* | -0.127 | 1.00 |
| *CAG.56* | -0.127 | 1.00 |
| *Ruminococcaceae_NK4A214_group* | -0.129 | 1.00 |
| *GCA.900066575* | -0.129 | 1.00 |
| *Ruminococcaceae_UCG.013* | -0.133 | 1.00 |
| *Lachnospiraceae_UCG.010* | -0.134 | 1.00 |
| *Lachnospiraceae_UCG.004* | -0.134 | 1.00 |
| *Ruminococcaceae_UCG.005* | -0.135 | 1.00 |
| *Roseburia* | -0.136 | 1.00 |
| *Lachnospiraceae_ND3007_group* | -0.142 | 1.00 |
| *Agathobacter* | -0.143 | 1.00 |
| *Ruminiclostridium_9* | -0.143 | 1.00 |
| *Oscillibacter* | -0.145 | 1.00 |
| *Ruminococcus_1* | -0.150 | 1.00 |
| *DTU089* | -0.161 | 1.00 |
| *Faecalibacterium* | -0.165 | 1.00 |
| *Subdoligranulum* | -0.165 | 1.00 |
| *Fusicatenibacter* | -0.167 | 1.00 |
| *Ruminococcus_2* | -0.172 | 1.00 |
| *Alistipes* | -0.173 | 1.00 |
| *Lachnospiraceae_FCS020_group* | -0.181 | 1.00 |
| *Ruminococcaceae_f* | -0.184 | 1.00 |
| *Butyricicoccus* | -0.186 | 1.00 |
| *Lachnospiraceae_NK4A136_group* | -0.193 | 1.00 |
| *Ruminococcaceae_UCG.002* | -0.203 | 1.00 |

**Supplementary Table 5**

**Bacterial counts in IgG2-sorted samples after removing reads found in technical controls**

| | Shared taxa | | |
|---|---|---|---|
| **Genus** | **Technical Controls** | **Samples** | **Post-correction sample counts** |
| *Blautia* | 33 | 6037 | 6004 |
| *Faecalibacterium* | 9 | 4180 | 4171 |
| *Pseudomonas* | 28 | 23 | 0 |
| *Aminobacter* | 841 | 36 | 0 |
| *Ruminiclostridium_5* | 32 | 613 | 581 |
| *Xanthobacteraceae_f* | 29 | 9 | 0 |
| *Bradyrhizobium* | 294 | 9 | 0 |
| *Coprococcus_3* | 1 | 1244 | 1243 |
| *Pseudomonas* | 95 | 23 | 0 |
| *Pseudomonas* | 55 | 39 | 0 |
| *Marvinbryantia* | 1 | 229 | 228 |
| *Blautia* | 3 | 17 | 14 |
| *Pseudomonas* | 10659 | 151 | 0 |
| *Lachnoclostridium* | 3 | 237 | 234 |
| *Blautia* | 2 | 45 | 43 |
| *Faecalibacterium* | 14 | 3549 | 3535 |
| *Streptococcus* | 4 | 19 | 15 |
| *Haemophilus* | 6 | 18 | 12 |
| *Faecalibacterium* | 3 | 108 | 105 |
| *Escherichia/Shigella* | 2 | 7259 | 7257 |
| *Corynebacterium_1* | 1 | 17 | 16 |
| *Sphingomonas* | 1545 | 8 | 0 |
| *Subdoligranulum* | 3 | 4456 | 4453 |
| *Acinetobacter* | 109 | 63 | 0 |
| *Micrococcus* | 39 | 100 | 61 |
| *Staphylococcus* | 62 | 91 | 29 |
| *Lachnospiraceae_f* | 1 | 78 | 77 |
| *Pseudomonas* | 30394 | 435 | 0 |
| *Anaerostipes* | 3 | 8715 | 8712 |
| *Blautia* | 2 | 69 | 67 |
| *Pseudomonas* | 3 | 50 | 47 |
| *Lachnospiraceae_ND3007_group* | 62 | 855 | 793 |
| *Mesorhizobium* | 2 | 10 | 8 |
| *Micrococcus* | 30 | 29 | 0 |
| *Achromobacter* | 1491 | 42 | 0 |
| *Lachnospiraceae_f* | 2 | 57 | 55 |
| *Coprococcus_2* | 1 | 49 | 48 |
| *Blautia* | 64 | 831 | 767 |
| *Coprococcus_2* | 14 | 1 | 0 |
| *Faecalibacterium* | 26 | 5497 | 5471 |
| *Faecalibacterium* | 21 | 6 | 0 |
| *Micrococcus* | 22 | 5 | 0 |
| *Fusicatenibacter* | 3 | 293 | 290 |
| *Staphylococcus* | 14 | 36 | 22 |

| | | | |
|---|---|---|---|
| *Blautia* | 26 | 1025 | 999 |
| *Ruminococcaceae_UCG-013* | 1 | 15 | 14 |
| *Mesorhizobium* | 50 | 4 | 0 |
| *Faecalibacterium* | 10 | 23 | 13 |
| *Faecalibacterium* | 17 | 5068 | 5051 |
| *Phascolarctobacterium* | 1 | 801 | 800 |
| *Corynebacterium_1* | 23 | 6 | 0 |
| *Stenotrophomonas* | 6969 | 157 | 0 |
| *Staphylococcus* | 15 | 4 | 0 |
| *Agathobacter* | 18 | 1 | 0 |
| *Enhydrobacter* | 167 | 38 | 0 |
| *Pseudomonas* | 375 | 297 | 0 |
| *Blautia* | 8 | 8 | 0 |
| *Mesorhizobium* | 146 | 118 | 0 |
| *Clostridium_sensu_stricto_1* | 71 | 35 | 0 |
| *Stenotrophomonas* | 3050 | 143 | 0 |
| *Pseudomonas* | 201 | 167 | 0 |
| *Achromobacter* | 1309 | 20 | 0 |
| *Ruminococcaceae_UCG-002* | 4 | 245 | 241 |
| *Mesorhizobium* | 8 | 11 | 3 |
| *Streptococcus* | 10 | 2923 | 2913 |
| *Blautia* | 24 | 10 | 0 |
| *Fusicatenibacter* | 4 | 5110 | 5106 |
| *Micrococcus* | 6 | 28 | 22 |
| *Blautia* | 10 | 213 | 203 |
| *Lachnospiraceae_ND3007_group* | 5 | 145 | 140 |
| *Mesorhizobium* | 1372 | 250 | 0 |
| *Lachnospiraceae_FCS020_group* | 2 | 129 | 127 |
| *Faecalibacterium* | 2 | 2 | 0 |
| *Ruminococcaceae_UCG-003* | 7 | 10 | 3 |
| *Lachnospiraceae_f* | 39 | 349 | 310 |
| *Aminobacter* | 251 | 8 | 0 |
| *Pseudomonas* | 24 | 6 | 0 |
| *Pseudomonas* | 133 | 134 | 1 |
| *Ruminiclostridium_5* | 11 | 10 | 0 |
| *Akkermansia* | 1 | 76 | 75 |
| *Lachnospiraceae_f* | 1 | 18 | 17 |
| *Pelomonas* | 42 | 66 | 24 |
| *Pseudomonas* | 46 | 53 | 7 |

#### Supplementary Table 6

**Mean relative abundance in sorted IgG2-coated vs. bulk samples**

| | | | IgG2-coated | | Bulk | |
|---|---|---|---|---|---|---|
| **Phylum** | **Family** | **Genus** | **Mean** | **SD** | **Mean** | **SD** |
| Firmicutes | *Acidaminococcaceae* | *Acidaminococcus* | 0.00E+00 | 0.00E+00 | 9.19E-05 | 1.65E-04 |
| Actinobacteria | *Actinomycetaceae* | *Actinomyces* | 0.00E+00 | 0.00E+00 | 7.17E-04 | 1.63E-03 |
| Firmicutes | *Lachnospiraceae* | *Agathobacter* | 1.27E-03 | 2.84E-03 | 2.85E-02 | 3.15E-02 |
| Verrucomicrobia | *Akkermansiaceae* | *Akkermansia* | 8.26E-04 | 1.44E-03 | 1.09E-03 | 2.79E-03 |
| Proteobacteria | *Burkholderiaceae* | *Alcaligenes* | 0.00E+00 | 0.00E+00 | 9.92E-05 | 2.16E-04 |
| Bacteroidetes | *Rikenellaceae* | *Alistipes* | 4.70E-04 | 9.53E-04 | 1.26E-02 | 2.17E-02 |
| Actinobacteria | *Bifidobacteriaceae* | *Alloscardovia* | 2.60E-04 | 6.88E-04 | 4.96E-05 | 1.31E-04 |
| Firmicutes | *Lachnospiraceae* | *Anaerobium* | 0.00E+00 | 0.00E+00 | 1.47E-03 | 3.90E-03 |
| Firmicutes | *Lachnospiraceae* | *Anaerosporobacter* | 0.00E+00 | 0.00E+00 | 5.41E-04 | 1.43E-03 |
| Firmicutes | *Lachnospiraceae* | *Anaerostipes* | 6.42E-02 | 8.55E-02 | 1.14E-02 | 1.38E-02 |
| Firmicutes | *Ruminococcaceae* | *Angelakisella* | 0.00E+00 | 0.00E+00 | 1.70E-04 | 4.50E-04 |
| Firmicutes | *Peptostreptococcaceae* | *Asaccharospora* | 1.21E-04 | 3.21E-04 | 1.75E-04 | 3.11E-04 |
| Actinobacteria | *Atopobiaceae* | *Atopobium* | 0.00E+00 | 0.00E+00 | 2.48E-04 | 6.57E-04 |
| Bacteroidetes | *Bacteroidaceae* | *Bacteroides* | 5.74E-02 | 1.48E-01 | 2.46E-01 | 2.15E-01 |
| Bacteroidetes | *Barnesiellaceae* | *Barnesiella* | 0.00E+00 | 0.00E+00 | 5.48E-03 | 1.45E-02 |
| Actinobacteria | *Bifidobacteriaceae* | *Bifidobacterium* | 7.89E-03 | 1.24E-02 | 1.30E-02 | 1.09E-02 |
| Proteobacteria | *Desulfovibrionaceae* | *Bilophila* | 2.14E-04 | 5.65E-04 | 5.46E-03 | 9.38E-03 |
| Firmicutes | *Lachnospiraceae* | *Blautia* | 1.30E-01 | 1.42E-01 | 2.91E-02 | 2.11E-02 |
| Firmicutes | *Ruminococcaceae* | *Butyricicoccus* | 1.09E-03 | 2.68E-03 | 1.51E-03 | 1.58E-03 |
| Bacteroidetes | *Marinifilaceae* | *Butyricimonas* | 0.00E+00 | 0.00E+00 | 3.61E-04 | 9.54E-04 |
| Firmicutes | *Ruminococcaceae* | *CAG-352* | 0.00E+00 | 0.00E+00 | 6.54E-06 | 1.73E-05 |
| Firmicutes | *Lachnospiraceae* | *CAG-56* | 1.18E-03 | 2.06E-03 | 8.10E-04 | 2.03E-03 |
| Epsilonbacteraeota | *Campylobacteraceae* | *Campylobacter* | 0.00E+00 | 0.00E+00 | 8.27E-06 | 2.19E-05 |
| Actinobacteria | *Eggerthellaceae* | *CHKCI002* | 0.00E+00 | 0.00E+00 | 9.44E-06 | 2.50E-05 |
| Firmicutes | *Christensenellaceae* | *Christensenellaceae_f* | 0.00E+00 | 0.00E+00 | 3.27E-05 | 8.66E-05 |
| Firmicutes | *Christensenellaceae* | *Christensenellaceae_R-7_group* | 2.51E-04 | 5.22E-04 | 6.51E-03 | 1.66E-02 |
| Firmicutes | *Clostridia_c* | *Clostridia_c* | 0.00E+00 | 0.00E+00 | 5.89E-05 | 1.56E-04 |
| Firmicutes | *Clostridiales_o* | *Clostridiales_o* | 2.89E-04 | 5.39E-04 | 8.89E-04 | 1.52E-03 |
| Firmicutes | *Clostridiales_vadinBB60_group* | *Clostridiales_vadinBB60_group_f* | 0.00E+00 | 0.00E+00 | 7.66E-04 | 2.03E-03 |
| Firmicutes | *Clostridiaceae_1* | *Clostridium_sensu_stricto_1* | 2.49E-02 | 5.10E-02 | 4.99E-02 | 8.48E-02 |
| Actinobacteria | *Coriobacteriaceae* | *Collinsella* | 0.00E+00 | 0.00E+00 | 6.72E-03 | 6.11E-03 |
| Bacteroidetes | *Barnesiellaceae* | *Coprobacter* | 0.00E+00 | 0.00E+00 | 1.75E-03 | 3.84E-03 |
| Firmicutes | *Lachnospiraceae* | *Coprococcus_1* | 5.82E-04 | 1.28E-03 | 6.23E-04 | 9.47E-04 |
| Firmicutes | *Lachnospiraceae* | *Coprococcus_2* | 2.02E-03 | 5.26E-03 | 3.42E-03 | 6.02E-03 |
| Firmicutes | *Lachnospiraceae* | *Coprococcus_3* | 5.13E-03 | 9.23E-03 | 1.69E-03 | 2.58E-03 |
| Actinobacteria | *Coriobacteriales_Incertae_Sedis* | *Coriobacteriales_Incertae_Sedis_f* | 0.00E+00 | 0.00E+00 | 3.27E-05 | 8.66E-05 |
| Firmicutes | *Defluviitaleaceae* | *Defluviitaleaceae_UCG-011* | 0.00E+00 | 0.00E+00 | 1.10E-04 | 2.36E-04 |
| Firmicutes | *Veillonellaceae* | *Dialister* | 1.63E-02 | 4.12E-02 | 1.85E-03 | 3.86E-03 |
| Firmicutes | *Lachnospiraceae* | *Dorea* | 1.66E-02 | 1.87E-02 | 5.70E-03 | 5.44E-03 |
| Firmicutes | *Ruminococcaceae* | *DTU089* | 2.08E-04 | 5.49E-04 | 1.48E-05 | 2.55E-05 |
| Actinobacteria | *Eggerthellaceae* | *Eggerthella* | 0.00E+00 | 0.00E+00 | 5.14E-05 | 1.36E-04 |
| Actinobacteria | *Eggerthellaceae* | *Eggerthellaceae_f* | 0.00E+00 | 0.00E+00 | 1.89E-05 | 5.00E-05 |
| Firmicutes | *Lachnospiraceae* | *Eisenbergiella* | 0.00E+00 | 0.00E+00 | 5.89E-05 | 1.56E-04 |
| Proteobacteria | *Enterobacteriaceae* | *Enterobacteriaceae_f* | 0.00E+00 | 0.00E+00 | 2.60E-03 | 6.89E-03 |
| Firmicutes | *Enterococcaceae* | *Enterococcus* | 2.37E-04 | 6.27E-04 | 6.02E-05 | 1.12E-04 |
| Firmicutes | *Lachnospiraceae* | *Epulopiscium* | 0.00E+00 | 0.00E+00 | 1.74E-04 | 4.60E-04 |
| Firmicutes | *Erysipelotrichaceae* | *Erysipelatoclostridium* | 0.00E+00 | 0.00E+00 | 2.69E-04 | 4.74E-04 |
| Firmicutes | *Erysipelotrichaceae* | *Erysipelotrichaceae_f* | 0.00E+00 | 0.00E+00 | 9.26E-05 | 2.45E-04 |
| Firmicutes | *Erysipelotrichaceae* | *Erysipelotrichaceae_UCG-003* | 4.18E-03 | 8.50E-03 | 1.96E-02 | 2.99E-02 |
| Proteobacteria | *Enterobacteriaceae* | *Escherichia/Shigella* | 6.92E-02 | 1.20E-01 | 5.33E-02 | 8.66E-02 |
| Firmicutes | *Ruminococcaceae* | *Faecalibacterium* | 1.06E-01 | 1.43E-01 | 3.39E-02 | 4.54E-02 |
| Firmicutes | *Erysipelotrichaceae* | *Faecalitalea* | 0.00E+00 | 0.00E+00 | 1.67E-04 | 4.04E-04 |
| Firmicutes | *Family_XIII* | *Family_XIII_AD3011_group* | 1.17E-04 | 3.10E-04 | 1.40E-04 | 3.43E-04 |
| Firmicutes | *Family_XIII* | *Family_XIII_f* | 0.00E+00 | 0.00E+00 | 2.62E-05 | 6.92E-05 |
| Firmicutes | *Family_XIII* | *Family_XIII_UCG-001* | 4.48E-05 | 1.18E-04 | 8.51E-05 | 2.25E-04 |

| | | | | | | |
|---|---|---|---|---|---|---|
| Firmicutes | *Family_XI* | *Finegoldia* | 0.00E+00 | 0.00E+00 | 7.21E-05 | 1.25E-04 |
| Firmicutes | *Ruminococcaceae* | *Flavonifractor* | 2.03E-04 | 5.38E-04 | 1.10E-03 | 2.22E-03 |
| Firmicutes | *Lachnospiraceae* | *Fusicatenibacter* | 3.14E-02 | 3.43E-02 | 8.23E-03 | 7.05E-03 |
| Fusobacteria | *Fusobacteriaceae* | *Fusobacterium* | 0.00E+00 | 0.00E+00 | 1.37E-04 | 1.85E-04 |
| Firmicutes | *Ruminococcaceae* | *GCA-900066225* | 0.00E+00 | 0.00E+00 | 1.31E-05 | 3.46E-05 |
| Firmicutes | *Lachnospiraceae* | *GCA-900066575* | 1.22E-04 | 3.24E-04 | 3.27E-05 | 8.66E-05 |
| Firmicutes | *Family_XI* | *Gemella* | 4.62E-04 | 1.22E-03 | 2.98E-04 | 7.88E-04 |
| Firmicutes | *Carnobacteriaceae* | *Granulicatella* | 3.25E-04 | 8.60E-04 | 5.30E-04 | 1.38E-03 |
| Proteobacteria | *Pasteurellaceae* | *Haemophilus* | 9.65E-04 | 1.72E-03 | 1.17E-02 | 2.27E-02 |
| Firmicutes | *Erysipelotrichaceae* | *Holdemanella* | 5.47E-04 | 1.45E-03 | 5.67E-04 | 1.50E-03 |
| Firmicutes | *Erysipelotrichaceae* | *Holdemania* | 0.00E+00 | 0.00E+00 | 3.27E-05 | 8.66E-05 |
| Firmicutes | *Lachnospiraceae* | *Hungatella* | 0.00E+00 | 0.00E+00 | 2.03E-04 | 3.43E-04 |
| Firmicutes | *Peptostreptococcaceae* | *Intestinibacter* | 3.39E-02 | 7.04E-02 | 3.51E-02 | 8.02E-02 |
| Firmicutes | *Ruminococcaceae* | *Intestinimonas* | 1.63E-04 | 2.43E-04 | 1.42E-04 | 2.49E-04 |
| Tenericutes | *Izimaplasmatales_o* | *Izimaplasmatales_o* | 0.00E+00 | 0.00E+00 | 5.23E-05 | 1.38E-04 |
| Proteobacteria | *Enterobacteriaceae* | *Klebsiella* | 5.86E-04 | 1.55E-03 | 3.88E-04 | 1.03E-03 |
| Firmicutes | *Lachnospiraceae* | *Lachnoclostridium* | 1.56E-02 | 1.84E-02 | 1.60E-02 | 1.51E-02 |
| Firmicutes | *Lachnospiraceae* | *Lachnospira* | 1.08E-02 | 1.56E-02 | 2.26E-02 | 1.96E-02 |
| Firmicutes | *Lachnospiraceae* | *Lachnospiraceae_AC2044_group* | 3.00E-04 | 7.94E-04 | 6.28E-05 | 1.07E-04 |
| Firmicutes | *Lachnospiraceae* | *Lachnospiraceae_f* | 7.90E-02 | 5.64E-02 | 2.38E-02 | 1.60E-02 |
| Firmicutes | *Lachnospiraceae* | *Lachnospiraceae_FCS020_group* | 2.87E-03 | 6.87E-03 | 3.69E-04 | 8.91E-04 |
| Firmicutes | *Lachnospiraceae* | *Lachnospiraceae_FE2018_group* | 0.00E+00 | 0.00E+00 | 3.91E-04 | 1.03E-03 |
| Firmicutes | *Lachnospiraceae* | *Lachnospiraceae_ND3007_group* | 7.24E-03 | 1.56E-02 | 2.56E-03 | 3.04E-03 |
| Firmicutes | *Lachnospiraceae* | *Lachnospiraceae_NK4A136_group* | 4.84E-03 | 7.26E-03 | 5.01E-03 | 5.98E-03 |
| Firmicutes | *Lachnospiraceae* | *Lachnospiraceae_UCG-001* | 6.93E-04 | 1.21E-03 | 5.78E-04 | 1.33E-03 |
| Firmicutes | *Lachnospiraceae* | *Lachnospiraceae_UCG-004* | 3.37E-03 | 4.55E-03 | 4.19E-03 | 3.25E-03 |
| Firmicutes | *Lachnospiraceae* | *Lachnospiraceae_UCG-010* | 2.15E-03 | 5.53E-03 | 7.80E-04 | 1.01E-03 |
| Firmicutes | *Lactobacillales_o* | *Lactobacillales_o* | 0.00E+00 | 0.00E+00 | 2.57E-04 | 6.80E-04 |
| Firmicutes | *Lactobacillaceae* | *Lactobacillus* | 9.57E-03 | 1.61E-02 | 5.58E-03 | 8.51E-03 |
| Firmicutes | *Streptococcaceae* | *Lactococcus* | 0.00E+00 | 0.00E+00 | 2.06E-05 | 5.44E-05 |
| Proteobacteria | *Pasteurellaceae* | *Mannheimia* | 0.00E+00 | 0.00E+00 | 4.10E-05 | 8.57E-05 |
| Firmicutes | *Lachnospiraceae* | *Marvinbryantia* | 3.42E-03 | 3.63E-03 | 2.60E-04 | 5.80E-04 |
| Firmicutes | *Veillonellaceae* | *Megamonas* | 0.00E+00 | 0.00E+00 | 4.24E-05 | 7.38E-05 |
| Firmicutes | *Veillonellaceae* | *Megasphaera* | 2.59E-04 | 4.03E-04 | 6.62E-05 | 1.75E-04 |
| Firmicutes | *Erysipelotrichaceae* | *Merdibacter* | 0.00E+00 | 0.00E+00 | 1.31E-05 | 3.46E-05 |
| Tenericutes | *Mollicutes_RF39_o* | *Mollicutes_RF39_o* | 0.00E+00 | 0.00E+00 | 8.25E-06 | 2.18E-05 |
| Proteobacteria | *Enterobacteriaceae* | *Morganella* | 0.00E+00 | 0.00E+00 | 9.32E-04 | 2.47E-03 |
| Bacteroidetes | *Muribaculaceae* | *Muribaculaceae_f* | 3.91E-04 | 1.03E-03 | 1.95E-03 | 5.17E-03 |
| Firmicutes | *Ruminococcaceae* | *Negativibacillus* | 0.00E+00 | 0.00E+00 | 2.22E-04 | 5.89E-04 |
| Bacteroidetes | *Marinifilaceae* | *Odoribacter* | 0.00E+00 | 0.00E+00 | 3.23E-03 | 4.10E-03 |
| Firmicutes | *Ruminococcaceae* | *Oscillibacter* | 4.17E-04 | 9.59E-04 | 1.36E-03 | 1.34E-03 |
| Firmicutes | *Ruminococcaceae* | *Oscillospira* | 5.18E-04 | 1.37E-03 | 3.14E-04 | 8.31E-04 |
| Bacteroidetes | *Tannerellaceae* | *Parabacteroides* | 7.06E-04 | 1.21E-03 | 1.33E-02 | 1.85E-02 |
| Bacteroidetes | *Prevotellaceae* | *Paraprevotella* | 0.00E+00 | 0.00E+00 | 2.64E-04 | 7.00E-04 |
| Proteobacteria | *Burkholderiaceae* | *Parasutterella* | 0.00E+00 | 0.00E+00 | 1.49E-02 | 2.74E-02 |
| Firmicutes | *Peptococcaceae* | *Peptococcaceae_f* | 3.41E-05 | 9.02E-05 | 1.24E-04 | 3.29E-04 |
| Firmicutes | *Family_XI* | *Peptoniphilus* | 6.25E-05 | 1.65E-04 | 2.71E-04 | 7.16E-04 |
| Firmicutes | *Peptostreptococcaceae* | *Peptostreptococcaceae_f* | 3.62E-05 | 9.59E-05 | 3.01E-05 | 7.95E-05 |
| Firmicutes | *Peptostreptococcaceae* | *Peptostreptococcus* | 6.99E-04 | 1.69E-03 | 1.80E-04 | 4.77E-04 |
| Firmicutes | *Acidaminococcaceae* | *Phascolarctobacterium* | 8.79E-03 | 2.31E-02 | 2.93E-03 | 7.72E-03 |
| Firmicutes | *Ruminococcaceae* | *Phocea* | 0.00E+00 | 0.00E+00 | 5.14E-05 | 1.36E-04 |
| Bacteroidetes | *Prevotellaceae* | *Prevotella* | 0.00E+00 | 0.00E+00 | 1.65E-05 | 4.37E-05 |
| Bacteroidetes | *Prevotellaceae* | *Prevotella_2* | 0.00E+00 | 0.00E+00 | 1.96E-05 | 5.19E-05 |
| Bacteroidetes | *Prevotellaceae* | *Prevotella_6* | 0.00E+00 | 0.00E+00 | 2.48E-05 | 6.55E-05 |
| Bacteroidetes | *Prevotellaceae* | *Prevotella_7* | 0.00E+00 | 0.00E+00 | 1.65E-05 | 4.37E-05 |
| Bacteroidetes | *Prevotellaceae* | *Prevotella_9* | 5.38E-03 | 1.42E-02 | 6.82E-02 | 1.78E-01 |
| Proteobacteria | *Enterobacteriaceae* | *Proteus* | 0.00E+00 | 0.00E+00 | 4.55E-04 | 1.06E-03 |
| Firmicutes | *Ruminococcaceae* | *Pseudoflavonifractor* | 0.00E+00 | 0.00E+00 | 1.96E-05 | 5.19E-05 |

| | | | | | | |
|---|---|---|---|---|---|---|
| Proteobacteria | *Rhodospirillales_o* | *Rhodospirillales_o* | 0.00E+00 | 0.00E+00 | 2.29E-03 | 5.98E-03 |
| Bacteroidetes | *Rikenellaceae* | *Rikenellaceae_RC9_gut_group* | 0.00E+00 | 0.00E+00 | 7.83E-05 | 1.53E-04 |
| Firmicutes | *Peptostreptococcaceae* | *Romboutsia* | 4.64E-02 | 1.17E-01 | 3.94E-02 | 1.03E-01 |
| Firmicutes | *Lachnospiraceae* | *Roseburia* | 8.55E-04 | 1.25E-03 | 2.43E-02 | 3.74E-02 |
| Proteobacteria | *Enterobacteriaceae* | *Rosenbergiella* | 0.00E+00 | 0.00E+00 | 2.06E-05 | 5.44E-05 |
| Actinobacteria | *Micrococcaceae* | *Rothia* | 2.82E-03 | 4.94E-03 | 1.16E-04 | 3.06E-04 |
| Firmicutes | *Ruminococcaceae* | *Ruminiclostridium_5* | 1.94E-03 | 4.14E-03 | 1.36E-03 | 1.85E-03 |
| Firmicutes | *Ruminococcaceae* | *Ruminiclostridium_6* | 0.00E+00 | 0.00E+00 | 6.01E-04 | 1.51E-03 |
| Firmicutes | *Ruminococcaceae* | *Ruminiclostridium_9* | 5.24E-04 | 1.06E-03 | 2.14E-03 | 2.75E-03 |
| Firmicutes | *Ruminococcaceae* | *Ruminococcaceae_f* | 2.01E-02 | 3.34E-02 | 2.37E-03 | 3.41E-03 |
| Firmicutes | *Ruminococcaceae* | *Ruminococcaceae_NK4A214_group* | 4.48E-03 | 9.09E-03 | 8.03E-04 | 1.47E-03 |
| Firmicutes | *Ruminococcaceae* | *Ruminococcaceae_UCG-002* | 7.75E-03 | 1.35E-02 | 8.25E-03 | 1.42E-02 |
| Firmicutes | *Ruminococcaceae* | *Ruminococcaceae_UCG-003* | 1.15E-04 | 3.03E-04 | 3.75E-03 | 7.40E-03 |
| Firmicutes | *Ruminococcaceae* | *Ruminococcaceae_UCG-004* | 0.00E+00 | 0.00E+00 | 9.16E-05 | 2.42E-04 |
| Firmicutes | *Ruminococcaceae* | *Ruminococcaceae_UCG-005* | 1.02E-03 | 1.69E-03 | 1.27E-03 | 3.32E-03 |
| Firmicutes | *Ruminococcaceae* | *Ruminococcaceae_UCG-009* | 0.00E+00 | 0.00E+00 | 7.20E-05 | 1.90E-04 |
| Firmicutes | *Ruminococcaceae* | *Ruminococcaceae_UCG-013* | 1.75E-03 | 2.44E-03 | 6.69E-04 | 1.59E-03 |
| Firmicutes | *Ruminococcaceae* | *Ruminococcaceae_UCG-014* | 4.89E-04 | 1.29E-03 | 6.71E-04 | 1.65E-03 |
| Firmicutes | *Ruminococcaceae* | *Ruminococcus_1* | 1.07E-02 | 2.83E-02 | 6.76E-03 | 1.44E-02 |
| Firmicutes | *Ruminococcaceae* | *Ruminococcus_2* | 1.61E-03 | 3.02E-03 | 6.72E-03 | 9.63E-03 |
| Bacteroidetes | *Marinifilaceae* | *Sanguibacteroides* | 0.00E+00 | 0.00E+00 | 3.73E-04 | 9.87E-04 |
| Firmicutes | *Lachnospiraceae* | *Sellimonas* | 1.58E-04 | 4.17E-04 | 3.93E-05 | 1.04E-04 |
| Actinobacteria | *Eggerthellaceae* | *Senegalimassilia* | 0.00E+00 | 0.00E+00 | 1.80E-04 | 4.77E-04 |
| Firmicutes | *Lachnospiraceae* | *Shuttleworthia* | 0.00E+00 | 0.00E+00 | 7.98E-05 | 1.59E-04 |
| Actinobacteria | *Eggerthellaceae* | *Slackia* | 0.00E+00 | 0.00E+00 | 1.65E-04 | 4.35E-04 |
| Firmicutes | *Erysipelotrichaceae* | *Solobacterium* | 0.00E+00 | 0.00E+00 | 8.27E-05 | 2.19E-04 |
| Firmicutes | *Streptococcaceae* | *Streptococcus* | 8.72E-02 | 1.66E-01 | 2.70E-02 | 5.91E-02 |
| Firmicutes | *Ruminococcaceae* | *Subdoligranulum* | 4.82E-02 | 1.11E-01 | 9.21E-03 | 1.62E-02 |
| Proteobacteria | *Burkholderiaceae* | *Sutterella* | 0.00E+00 | 0.00E+00 | 1.40E-02 | 2.65E-02 |
| Firmicutes | *Peptostreptococcaceae* | *Terrisporobacter* | 2.84E-03 | 7.50E-03 | 1.70E-03 | 4.50E-03 |
| Firmicutes | *Erysipelotrichaceae* | *Turicibacter* | 5.42E-04 | 1.42E-03 | 4.63E-03 | 8.07E-03 |
| Firmicutes | *Lachnospiraceae* | *Tyzzerella* | 0.00E+00 | 0.00E+00 | 4.45E-04 | 8.41E-04 |
| Firmicutes | *Lachnospiraceae* | *Tyzzerella_3* | 9.66E-04 | 1.85E-03 | 9.01E-04 | 2.21E-03 |
| Firmicutes | *Lachnospiraceae* | *Tyzzerella_4* | 6.58E-03 | 1.17E-02 | 1.78E-03 | 3.17E-03 |
| Firmicutes | *Ruminococcaceae* | *UBA1819* | 7.19E-05 | 1.70E-04 | 1.71E-04 | 2.52E-04 |
| Firmicutes | *Lachnospiraceae* | *UC5-1-2E3* | 0.00E+00 | 0.00E+00 | 2.62E-05 | 6.92E-05 |
| Firmicutes | *Veillonellaceae* | *Veillonella* | 5.39E-03 | 6.91E-03 | 3.72E-02 | 8.55E-02 |
| Firmicutes | *Veillonellaceae* | *Veillonellaceae_f* | 1.53E-04 | 4.06E-04 | 2.65E-04 | 4.30E-04 |
| Lentisphaerae | *Victivallaceae* | *Victivallis* | 0.00E+00 | 0.00E+00 | 1.96E-05 | 5.19E-05 |
| Firmicutes | *Leuconostocaceae* | *Weissella* | 1.20E-03 | 3.13E-03 | 5.46E-04 | 1.44E-03 |
