## Supplementary Figures for "Specific gut pathobionts escape antibody coating and are enriched during flares in patients with severe Crohn’s disease"

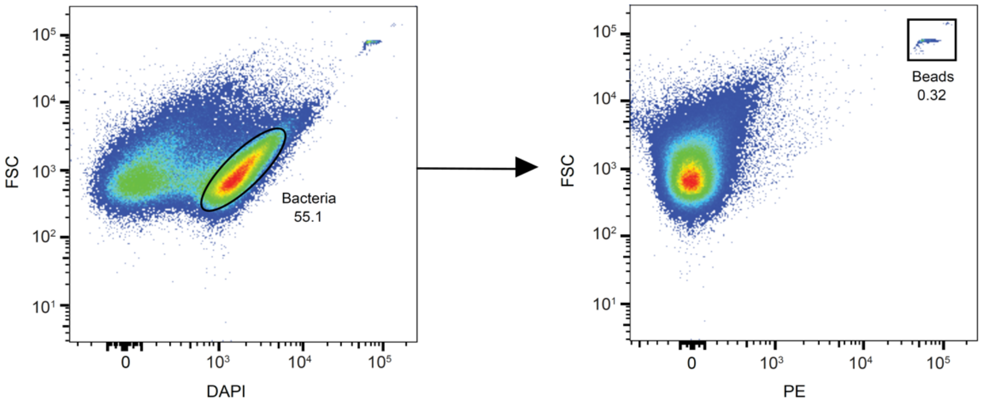


**Supplementary Figure 1. Quantification of gut bacteria in stool by flow cytometry.** Single cell suspensions of stool bacteria were labelled with DAPI to exclude debris (DAPI gate on the left), and then counted based on gating of count beads added to the same tube (right).

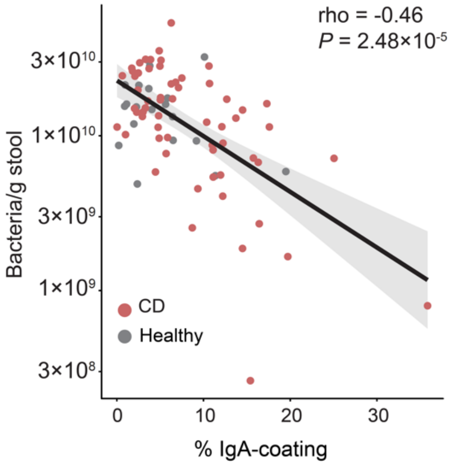


**Supplementary Figure 2. % IgA-coating is inversely related to gut bacterial load.** Scatter plot showing the correlation between % IgA-coated bacteria and bacteria/g stool in CD patients (n=60) and healthy controls (n=20). Spearman's rho statistics was used for correlations. The line and shaded area depict a linear fit, and the 95% confidence interval, respectively.

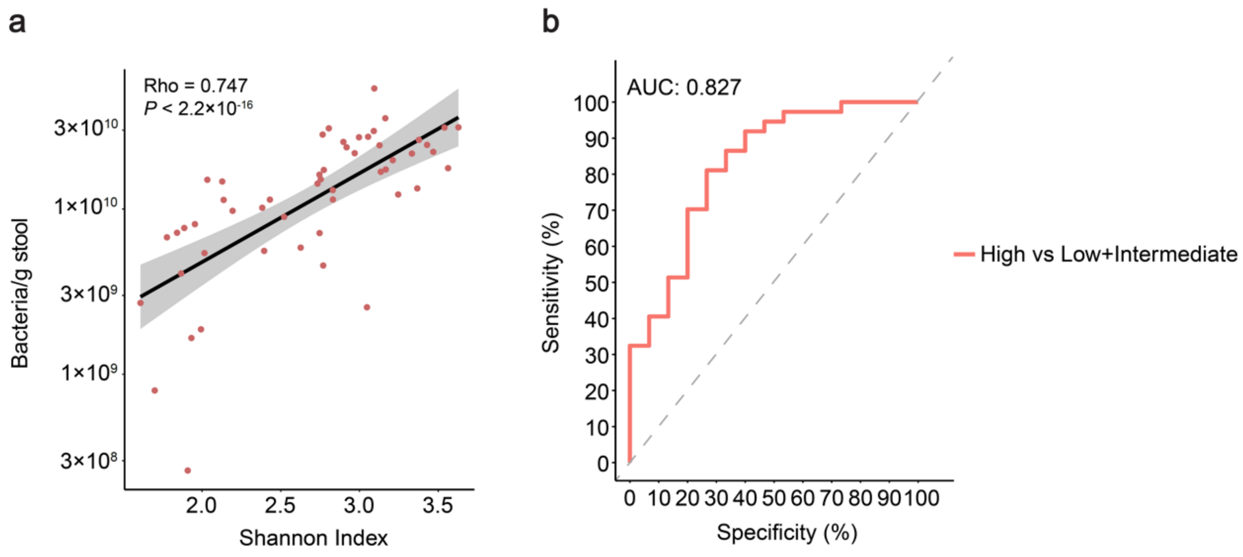


**Supplementary Figure 3. Shannon index in relation to bacteria/g stool in CD patients and sPLS-DA ROC curve.** a) Scatterplot of Shannon index in relation to number of bacteria/g stool in CD patients (n=52). Statistical test and plotting details are described in Supplementary Figure 2. b) ROC curve showing the AUC for the sPLS-DA analysis used to identify IgG2-hi indicator genera.

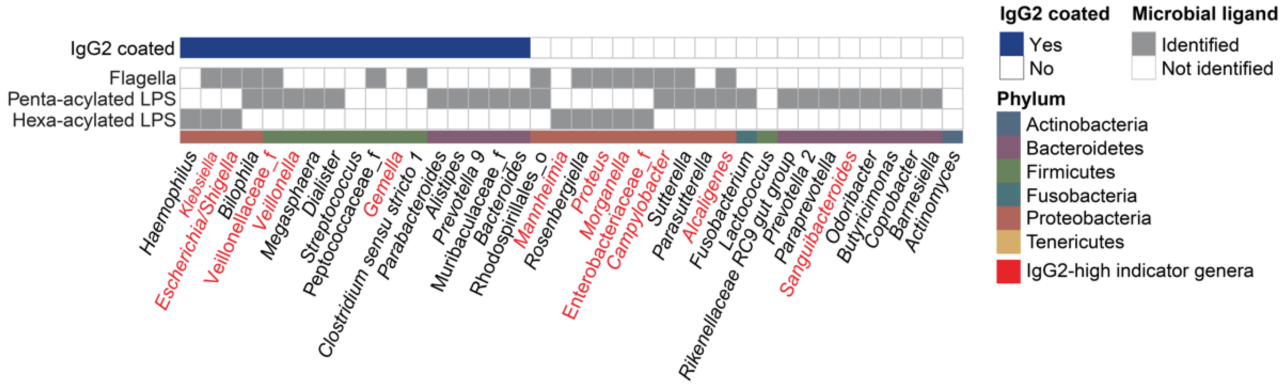


**Supplementary Figure 4. Selected innate immune activating ligands in IgG2-coated and non-coated bacteria.** Lower panel shows genome-based identification of innate immune activating ligands (grey squares) in bacteria identified in Figure 3 using PICRUST2 analysis. Bacteria found to be IgG2-coated are marked with a blue square in the upper panel. Bacteria highlighted in red represent IgG2-hi indicator genera.

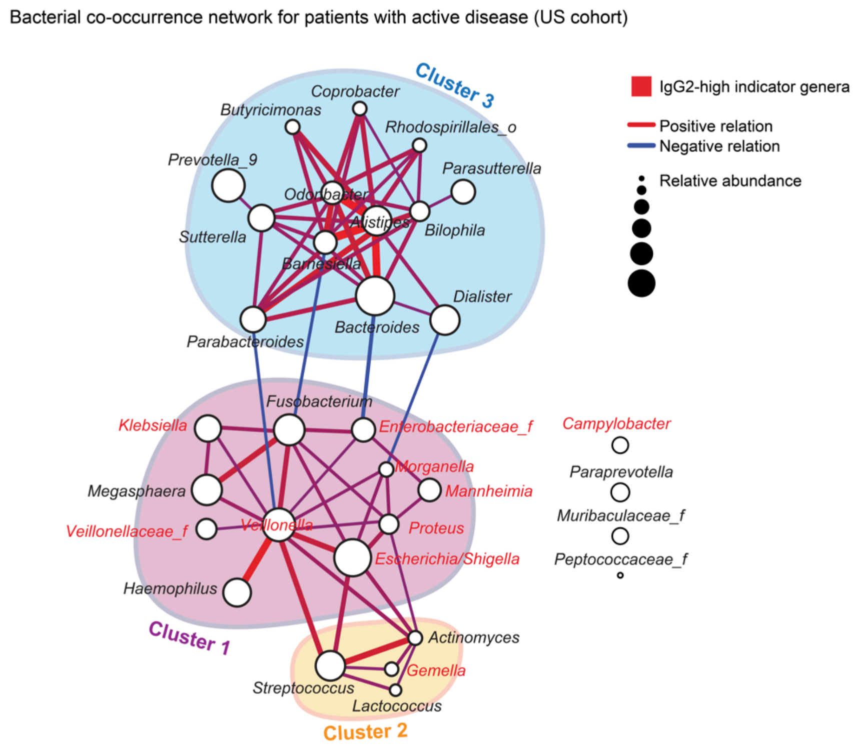


**Supplementary Figure 5. Bacterial co-occurrence network in CD patients with active disease from US cohort.** Co-occurrence network of bacteria from Figure 5a in CD patients with active disease (n=205) from the US cohort. Clusters were identified using the walktrap algorithm and are indicated by an orange, blue or purple circle. Red and blue edges represent positive and negative relations, respectively. Node sizes represent relative bacterial abundance.
